# A Pan-Cancer Single-Cell Compendium of Intratumoural Heterogeneity

**DOI:** 10.64898/2026.01.06.693992

**Authors:** Ido Nofech-Mozes, Ashton Cook, Tom W Ouellette, Sara Hafezi-Bakhtiari, Philip Awadalla, Sagi Abelson

## Abstract

Intratumoural heterogeneity remains one of the most formidable challenges in oncology, driving treatment resistance, disease progression, and poor patient outcomes. To uncover the cellular programs that define tumour composition and diversity, we interrogated over 3.6 million single cells derived from more than 1,000 primary and metastatic tumours spanning 14 diverse cancer types. Through the identification and quantification of coordinated gene expression programs, we delineated heterogeneous cancer cell states, exhibiting unique and clinically significant associations with patient survival. Integration with high-resolution spatial transcriptomics identified prognostic cancer cell states marked by aggressive transcriptional programs, co-localizing with invasive histological features and significantly enriched in metastatic settings. Functional drug perturbation screening uncovered drug vulnerabilities specific to distinct cancer cell states, supporting both direct targeting strategies and therapeutic reprogramming toward less aggressive cellular phenotypes. Collectively, this study offers novel insights into the relationship between intratumoural transcriptional heterogeneity, spatial organization, clinical outcomes, and therapeutic vulnerabilities.

---

Intratumoural heterogeneity (ITH) is a defining characteristic of human cancers, reflecting the presence of malignant subpopulations with distinct molecular and functional properties. This heterogeneity arises from a combination of genomic alterations, epigenetic regulation, and transcriptional plasticity, enabling tumours to adapt to diverse selective pressures^1,2^. ITH contributes to core hallmarks of cancer such as sustained proliferation, immune evasion, and metastatic progression, and represents a major barrier to durable clinical responses^3–7^. Understanding the mechanisms that generate, sustain, and exacerbate ITH is essential for improving cancer diagnosis, prognosis, and treatment^8^.

In aggressive solid tumours, single-cell analyses have revealed profound cellular heterogeneity. For example, in glioblastoma, malignant cells have been shown to transition between transcriptional states that reflect neural development programs, with these shifts contributing to treatment resistance and disease progression^9^. In pancreatic ductal adenocarcinoma, single-cell transcriptomic analysis has revealed malignant subpopulations occupying distinct transcriptional states that differ in differentiation status, proliferative capacity, and clinical outcomes^10^. Together, these observations underscore the biological importance of transcriptional cell states, demonstrating that malignant cells within a tumour can adopt diverse, functionally meaningful programs that shape tumour behavior.

Despite these insights, defining stable and reproducible transcriptional states remains challenging, in part because cell states arise from complex gene-expression programs that can be parsed at multiple biological and analytical levels. For example, two malignant subpopulations within the same tumour may engage distinct cell-cycle gene modules, yet both are classified broadly as proliferative^2^. While both cancer-type-specific^7,11,12^ and pan-cancer^13,14^ studies have characterized transcriptional heterogeneity in malignant cells, multiple methodological frameworks exist. This complexity is further compounded by the co-expression of multiple programs within individual cells, producing a combinatorial diversity of transcriptional subpopulations that is challenging to interpret. Moreover, transcriptional divergence does not always translate into meaningful clinical outcomes, highlighting the need for systematic evaluation to identify which transcriptional states carry biological or prognostic significance

In this study, we systematically define distinct gene expression programs of malignant subpopulations within and across common and clinically challenging cancer types and elucidate their spatial organization, prognostic significance, and therapeutic vulnerabilities, advancing both the mechanistic understanding and translational relevance of intratumoural heterogeneity.

## Results

### Constructing a pan-cancer single-cell atlas for characterizing tumour heterogeneity

To define cancer cell states across solid tumours, we curated a large single-cell RNA sequencing (scRNA-seq) atlas from publicly available datasets (**Supplementary Table 1**). This compendium includes 1,355 patient-derived samples from 38 studies, comprising 1,021 tumours across 14 cancer types, as well as 334 matched normal and tumour-adjacent biopsies. Clinical annotations, including age, sex, tumour stage, treatment status, and histological grade, are available for most samples and integrated into the pan-cancer atlas (**Figure 1a-b**).

**Figure 1:**
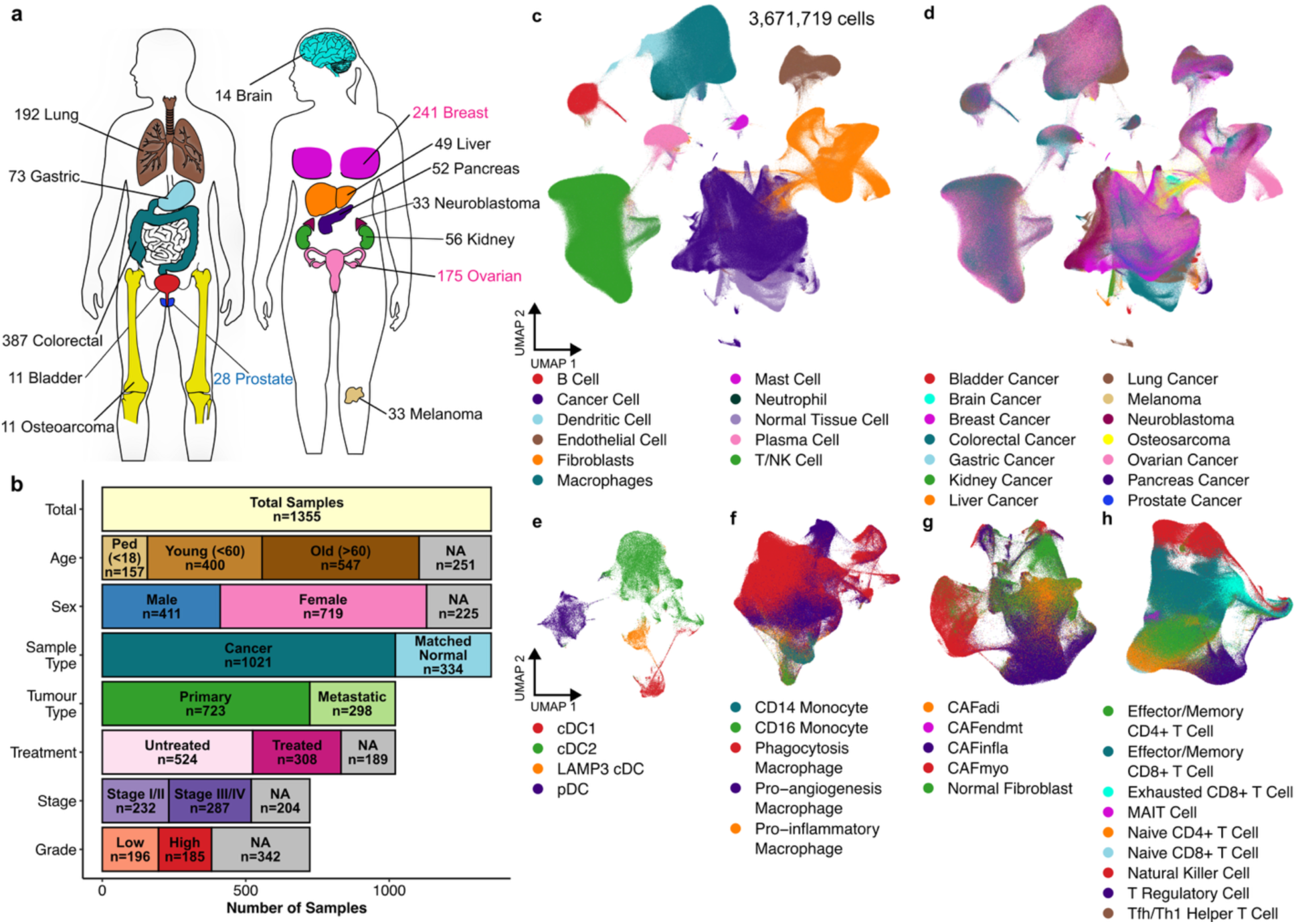
Overview of the curated pan-cancer single-cell atlas. **a,** Distribution of samples by tissue of origin. Male and female specific cancer types are indicated by blue and pink text respectively. **b,** Clinical characteristics of the samples in the atlas. NA denotes missing clinical data. **c-d,** UMAP visualization of the integrated pan-cancer atlas coloured by major cell types (**c**) and tissue of origin of the samples (**d**). **e-h,** UMAP visualization of granular cell subtypes in the tumour microenvironment, identified by scATOMIC and annotated as dendritic cells (**e**), macrophages (**f**), fibroblasts (**g**), and T/NK cells (**h**).

To ensure consistency, we harmonized cell type annotations across a total of 3,671,719 cells using our previously developed classification framework, scATOMIC^15^ (**Figure 1c–h**). To distinguish malignant cells from normal tissue cells, we employed a stringent two-step approach. First, cells were scored for a malignant gene expression signature using scATOMIC^15^ on a per-patient basis. Second, copy number variation (CNV) profiles were independently inferred using CopyKAT (RRID:SCR_024512)^16^. Only cells consistently classified as malignant by both methods (n = 737,791) were retained for downstream analyses of cancer cell heterogeneity. Across all cancer types, scATOMIC and CopyKAT exhibit a high level of concordance in malignant cell detection, with each method identifying about 74% of the malignant cells detected by the other. (**Supplementary Figure 1**). Each approach also detected unique malignant cell subsets, underscoring the value of integrating detection methods to achieve high confidence. Melanoma samples were omitted from downstream analysis as only a small number of cancer cells showed concordant scATOMIC and CopyKAT predictions.

To support transparency and reuse, we made the full compendium publicly available as a unified AnnData object^17,18^ (**Data Availability**).

### Identification of coordinated gene modules driving tumour heterogeneity

Gene-gene covariance analysis provides an unsupervised framework for identifying functional gene programs within cell types^19–21^. To prioritize covariance patterns that reflect biologically meaningful variation, both within and across patient biopsies of a given tumour type, we applied Hotspot, a graph-based method that identifies gene modules exhibiting structured expression patterns across the transcriptional landscape^22^. Analyses were performed on integrated scVI latent representations, incorporating sample identifier as a batch variable^23^. This approach yielded 351 gene expression modules across the 13 cancer types analyzed, with an average of 27 modules per cancer type.

Hotspot modules revealed substantial presence of coordinated gene programs within individual cancer types. For example, in lung adenocarcinoma (LUAD, **Figure 2a**), Hotspot identified 33 gene modules, many of which showed strong pairwise correlations in gene-level Z-scores, suggesting that certain modules share common gene components (**Figure 2b**). This overlap contributed to redundancy at the single-cell level, where individual malignant cells frequently co-expressed multiple related modules, while also reflecting the combinatorial nature of transcriptional programs in tumours (**Supplementary Figure 2**).

**Figure 2:**
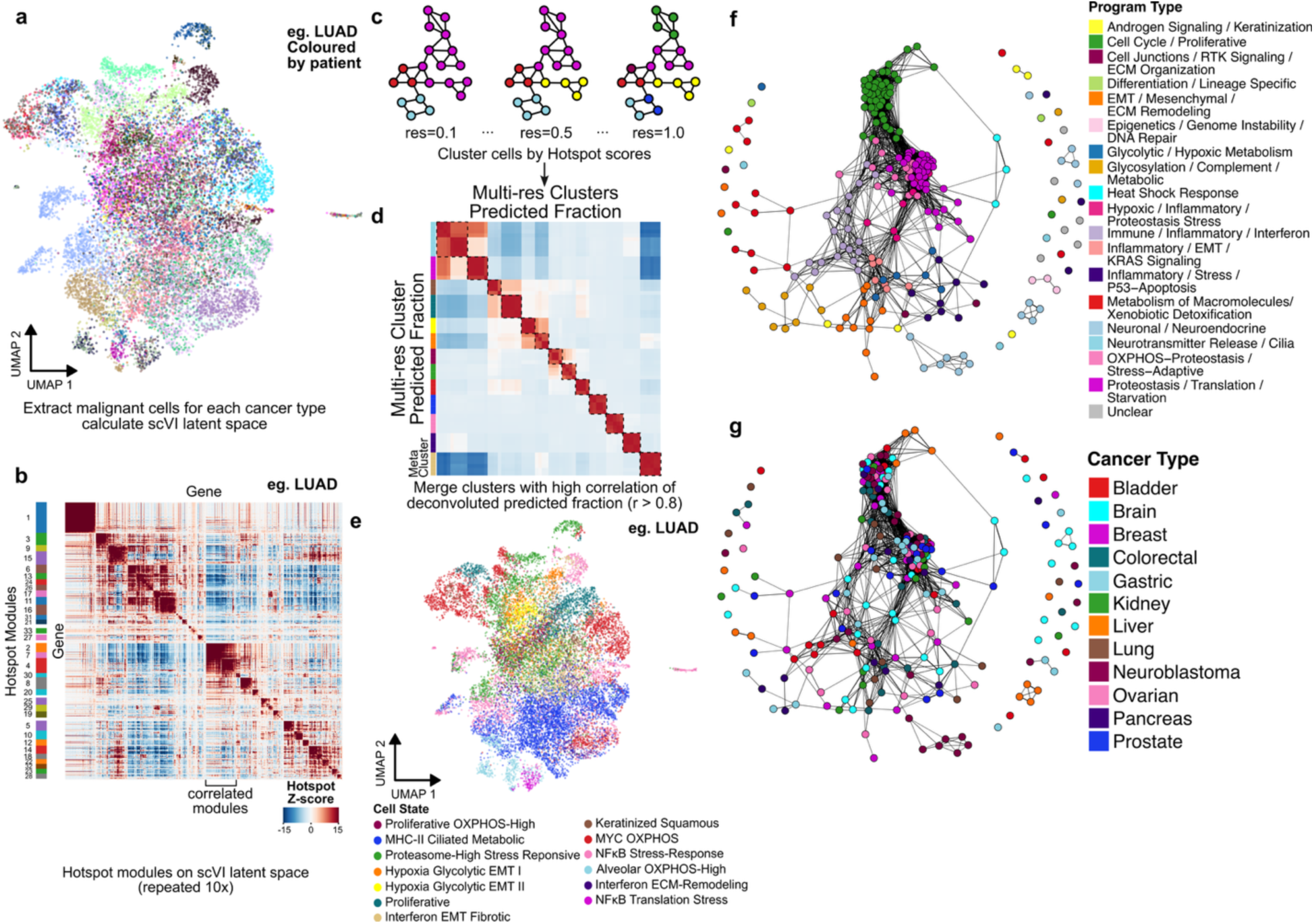
Defining cancer cell states through clustering of Hotspot module scores and deconvolution of bulk gene expression cohorts. **a**, UMAP visualization of the scVI-derived latent space for 20,654 lung adenocarcinoma cells from 62 primary tumour samples, coloured by patient. **b**, Heatmap showing gene-gene covariance in LUAD, with colours representing Hotspot-derived Z-scored correlations. Hotspot modules are labeled on the y-axis and modules 2, 4, and 7 are highlighted to illustrate highly correlated modules. **c,** Cells were clustered using the Louvain method across 10 resolutions (0.1–10). A schematic of three representative resolutions is shown, with cells coloured according to their cluster assignments. **d**, Heatmap showing Pearson correlation values between predicted deconvoluted proportions of each cell state across clustering resolutions. Louvain clusters at multiple resolutions were merged into meta-clusters based on correlated predicted fractions (>0.8). Results from the LUAD cohort are presented as an illustrative example. **e**, UMAP visualization of the LUAD dataset, with cells coloured according to their assigned cell state. **f-g**, Pan-cancer cell state similarity network. Nodes represent distinct cell states across the pan cancer single-cell atlas coloured by their respective core programs (**f**) and cancer type (**g**). The proximity of nodes represents the p-value-weighted Jaccard similarity between upregulated pathways in each cell state.

### Defining discrete transcriptional states underlying intratumoural heterogeneity

Defining discrete transcriptional states offers a practical approach for resolving overlapping gene programs, enhancing biological interpretability, and facilitating the study of ITH. To resolve discrete transcriptomic states, we clustered malignant cells based on their Hotspot-derived gene module activity profiles, which reflect the collective expression strength of each module’s genes in individual cells^22^. For each cancer type, we constructed a nearest-neighbor graph of Hotspot module scores and applied Louvain community detection across a range of resolution settings, capturing both broad cellular phenotypes and finer transcriptional distinctions (**Figure 2c**).

To prioritize cell states with translational potential, we next evaluated whether these transcriptomic identities could be reliably detected in bulk RNA-seq data. Using BayesPrism^24^, we deconvoluted primary tumour samples from the TCGA^25^ and TARGET^26^ cohorts, to estimate the relative abundance of each single-cell-defined cancer cell state.

Given that cell states identified at different clustering resolutions may overlap, we correlated the deconvoluted cancer cell state proportions between resolutions and across patients. Highly correlated clusters (Pearson’s r > 0.8) were merged into meta-clusters to refine the classification (**Figure 2d; Methods**). This approach identified an average of 18.5 cell states per cancer type. At the patient level, each tumour sample had a median of four distinct cell states occurring at >5% cell fraction, highlighting both inter-patient heterogeneity and the tendency of individual tumours to be dominated by a smaller subset of states.

To better interpret the biological functions associated with the identified cancer cell states, we performed gene set enrichment analysis on genes found to be upregulated within each meta-cluster compared to all others^27^. These revealed pathways reflecting distinct cellular functions (**Supplementary Table 2**). For example, in LUAD, cell states were characterized by processes including cell cycle, hypoxia response, and protein regulation (**Supplementary Figure 3**). Unique functional labels were then assigned to each cancer cell in the pan-cancer atlas based on the pathway enrichment profile of its corresponding cell state (**Figure 2e, Supplementary Table 2**).

Lastly, a pan-cancer similarity network of cell states was constructed (**Methods**), revealing recurrent, generalized programs, including those related to cell cycle, metabolism, protein regulation, interferon response, and extracellular matrix remodeling, that are shared across multiple cancer types (**Figure 2f, g**). Conversely, some cell states exhibited tissue-specific signatures, such as neuronal and neuroendocrine programs enriched in glioblastoma and neuroblastoma.

To support future adoption and reproducible identification of cancer cell states in scRNA-seq data, we developed ITHmapper, an open-source Python package that enables robust annotation of both cancer-type cell states and pan-cancer generalized programs in new, query single-cell datasets (**Supplementary Figure 4**).

### Cell state abundance serves as a functional biomarker linking tumour heterogeneity to clinical prognosis

The presence of different malignant cell populations within a tumour reflect its underlying biological complexity and clinical behavior^24,28^. To assess the prognostic value of the identified cancer cell states, we revisited their estimated abundance across patient samples using deconvolution of bulk RNA-seq data. (**Figure 3a**, **Supplementary Table 3**). Prognostic significance was evaluated by determining optimal quantile-based thresholds to stratify patients into high- and low-abundance groups within discovery cohorts (TCGA^25^ for adult cancers and TARGET^26^ for neuroblastoma), followed by validation in independent datasets (**Extended Data Figure 1, Supplementary Table 4**).

**Figure 3:**
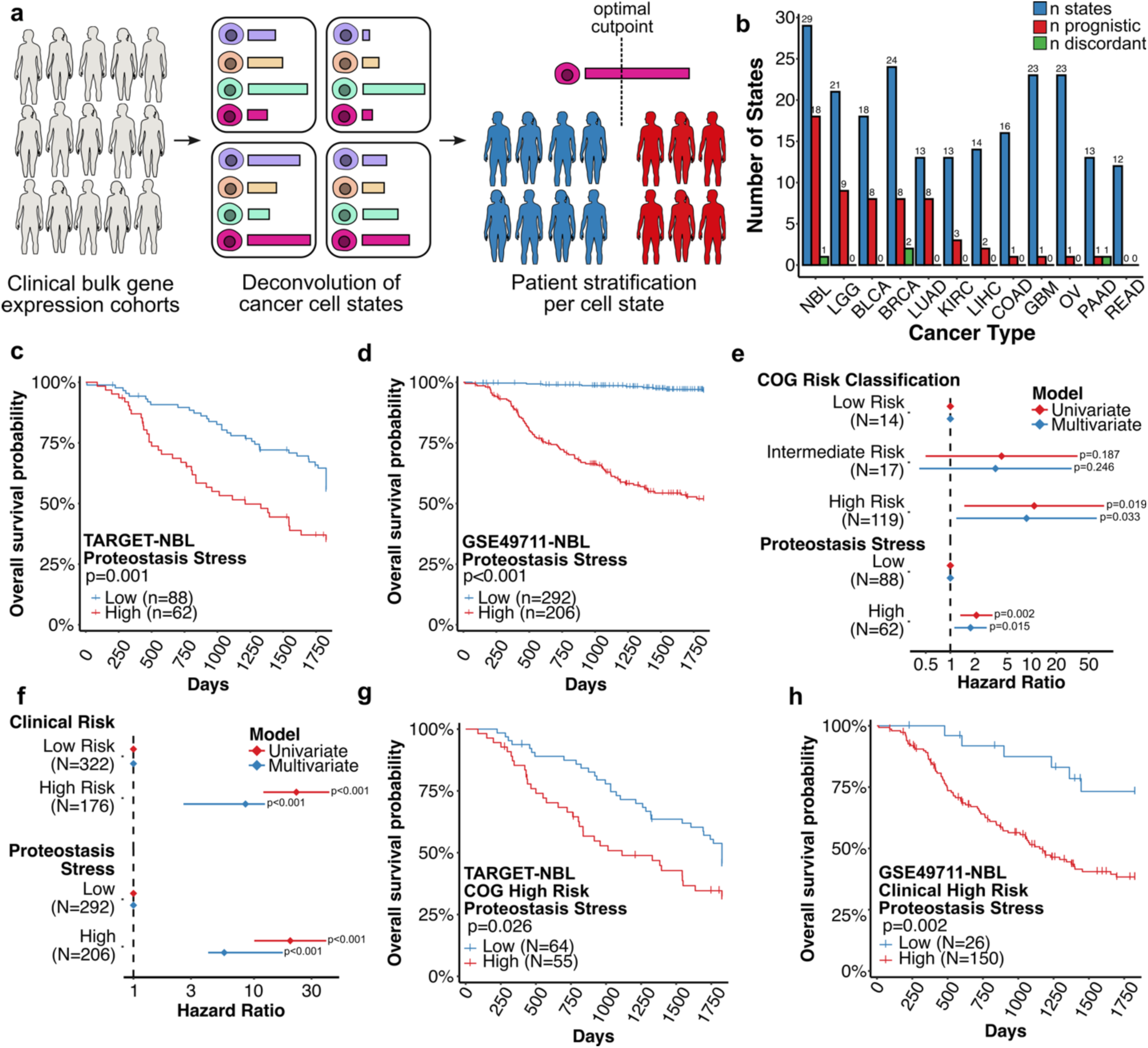
Cell state deconvolution identifies prognostic population in neuroblastoma independent from clinical risk. **a**, Schematic of the deconvolution and patient stratification workflow, indicating how patients are categorized into high and low-abundance groups based on inferred cell state proportions. Patients are stratified independently for each cell state. **b**, Barplot depicting the number of cancer cell states per cancer type that were found to be prognostic or discordant between the discovery and validation cohorts. Cell states were considered as prognostic if their hazard ratios had a P < 0.05 in both datasets and exhibited concordant directional associations with survival. Cell states were defined as discordant if their hazard ratios had a P < 0.05 in discovery and validation datasets with opposite directional survival associations. **c**,**d**, Kaplan-Meier plots for the *Proteostasis Stress* neuroblastoma cell state in TARGET (**c**) and the GSE49711 NBL cohorts (**d**). High-abundance patients represent those with elevated inferred proportions of the *Proteostasis Stress* cell state, while low-abundance patients have comparatively reduced representation of this program. **e**-**f,** Univariate and multivariate hazard ratios with 95% confidence intervals are show for clinical risk classification and patient stratification based on the proportion of *Proteostasis Stress* neuroblastoma cells in TARGET-NBL (**e**) and in GSE49711 (**f**). **g**-**h**, Kaplan–Meier curves based on *Proteostasis Stress* cell-state abundance in high-risk neuroblastoma, defined by COG criteria in TARGET-NBL (**g**) and by clinical high-risk classification in GSE49711 (**h**). Patients with high cell-state abundance are shown in red, and those with low abundance in blue. Significance was assessed using the log-rank test.

The proportion of cancer cell states with prognostic potential varied across cancers (**Figure 3b**). For example, in neuroblastoma (NBL), 18 of 29 cell states (62%) were significantly associated with survival rates (log rank test; P < 0.05), whereas no significant associations were observed in rectal adenocarcinoma (READ). This variability among cancers partly reflects differences in cohort characteristics, including sample size, sequencing or array technology, patient heterogeneity within and between the discovery and validation cohorts, and outcome distribution. The impacts of cohort or experimental design are particularly pronounced among cancers with rapid post-diagnosis mortality, such as pancreatic cancer and glioblastoma, where limited variability in survival outcomes reduces statistical power to detect prognostic associations (**Supplementary Table 4**).

The variability in prognostic associations across cancer types prompted a more focused analysis in NBL, where multiple cell states showed a strong effect on survival (hazard ratio > 2). Despite advances in molecular profiling and risk stratification, outcome prediction within high-risk neuroblastoma patients remains challenging^29^. The Children’s Oncoalogy Group (COG) risk classification system^30^, which is routinely used in clinical practice, integrates multiple variables, including patient age at diagnosis, disease stage, tumour histology, MYCN amplification status, chromosomal aberrations (e.g., 1p/11q loss, 17q gain), and DNA ploidy. To determine whether transcriptionally defined cell states provide additional prognostic information beyond these clinical factors, we examined the *Proteostasis Stress* cell state in NBL as it exhibited strong significant associations with patient survival in both the discovery and validation cohorts^26,31^ (**Figure 3c,d**). Notably, this association remained significant after adjusting for the COG risk score reported in the discovery cohort using a multivariate Cox model (HR = 1.8, P < 0.05; **Figure 3e**).

In the validation cohort, where clinically high-risk patients were defined as stage 4 disease diagnosed after 18 months of age or by the presence of MYCN amplification regardless of age or stage^31^, the *Proteostasis Stress* cell state similarly stratified patients into distinct prognostic subgroups, independent of this clinical classification (HR = 5.7, P < 0.001,**Figure 3f)**. Among patients clinically classified as high-risk in both datasets, the *Proteostasis Stress* cell state further stratified individuals into risk groups and remained prognostic (**Figure 3g-h**; log-rank P = 0.026 and P = 0.002, respectively).

Together, these findings demonstrate that cancer cell states defined by single-cell transcriptomics provide prognostic information that can complement and extend established clinical risk classifications.

### Spatial mapping of cancer cell states reveals distinct architectural patterns linked to histology, invasion, and genomic features

To examine malignant cell states in their spatial context within tumours, we analyzed high-resolution public Visium HD transcriptomic data from LUAD (termed here as Patient 1), colorectal adenocarcinoma (COAD; Patients 2-4), and breast cancer (BRCA; Patients 5-6)^32–35^. The Visium HD platform enables whole-transcriptome profiling at subcellular (2 × 2 µm) resolution, allowing direct mapping of cancer cell states onto spatially resolved single cells^36^. Cell type annotation was performed using a combination of scATOMIC^15^ and manual curation, and cancer type–specific cell states were assigned to malignant cells using ITHmapper (**Methods**, **Extended Data Figure 2, Supplementary Figure 4**).

In LUAD, we observed three distinct histological growth patterns: papillary, micropapillary, and acinar (**Figure 4a**). Poor-prognosis cell states identified in Patient 1, including *Proteasome-High Stress Responsive* and *Hypoxia Glycolytic EMT I* localized preferentially to the papillary and micropapillary regions, while cell states associated with more favorable outcomes, including *MHC-II Ciliated Metabolic* and *NFκB Stress Responsive*, were enriched in the acinar areas (**Figure 4a-b, Extended Data Figure 3**).

**Figure 4:**
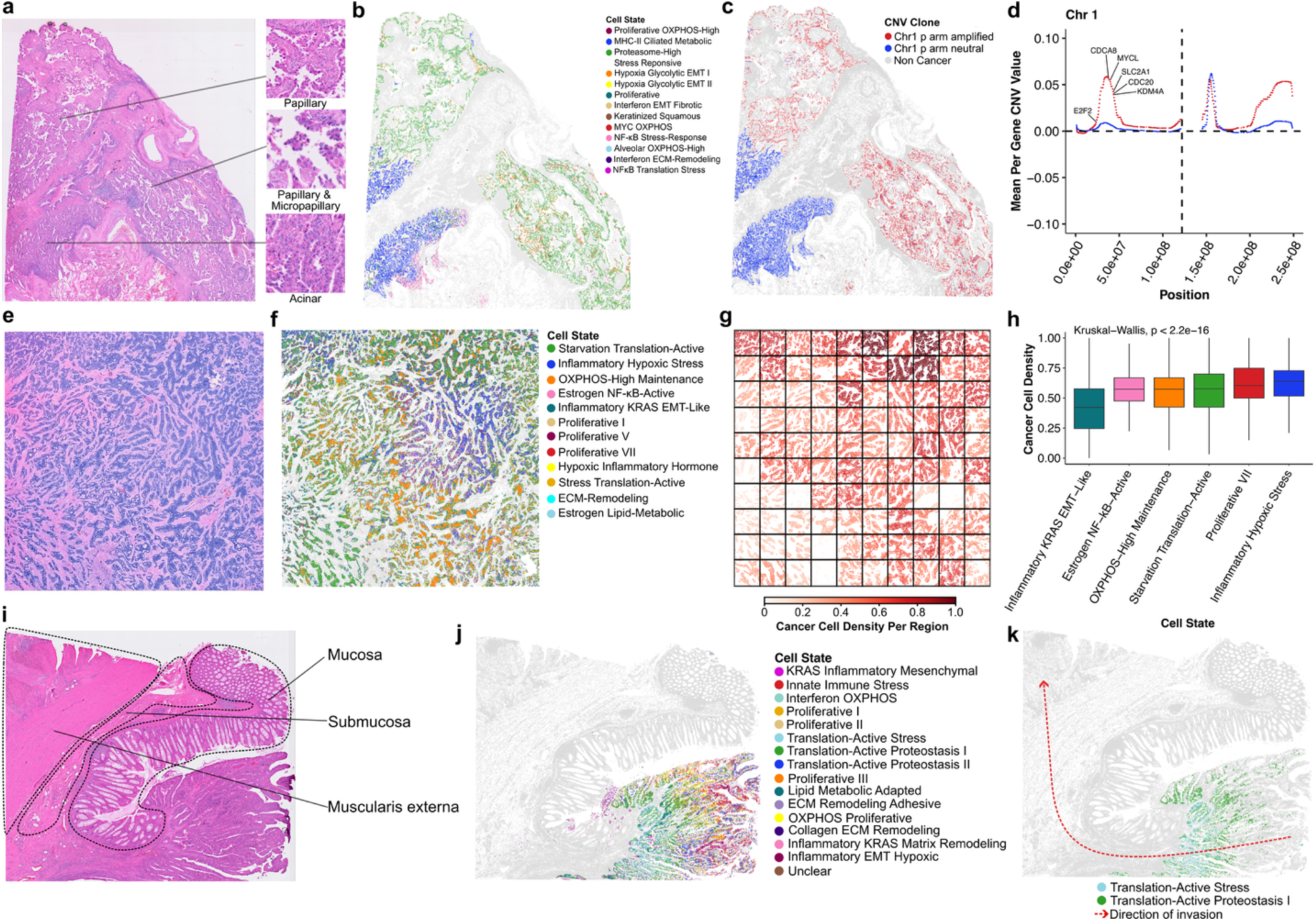
Spatial localization patterns of cancer cell states. **a,** Hematoxylin and eosin (H&E) staining of LUAD Patient 1. Regions of variable histology growth patterns are highlighted. **b-c,** Spatial representation of LUAD cells coloured by mapped cancer cell state **(b)** and inferred CNV profile cluster **(c). d,** Inferred CNV values along chromosome 1, showing average values per gene for cells classified as chr1 p-arm amplified or chr1 p-arm neutral. The dashed line indicates the genomic position of the centromere. **e,** H&E staining of BRCA Patient 3. **f-g,** spatial representation of BRCA cells coloured by mapped cancer cell state **(f)** and scaled cancer cell density **(g). h,** Boxplot depicting scaled cancer cell densities across cancer cell states identified in the tumour section. Values represent the cancer cell density within the region plotted in (**g**) for each cell of a respective cell state. Boxes and whiskers represent the lower fence, first quartile (Q1), median (Q2), third quartile (Q3), and upper fence. Significance was calculated using a Kruskal-Wallis test. **i,** H&E staining of COAD Patient 4. Dotted lines represent the layers of the normal epithelium. **j,** Spatial representation of COAD cells coloured by their corresponding cancer cell state. **k,** Enrichment of *Translation-Active Stress* and *Translation-Active Proteostasis I* COAD cells in invasive front of the tumour.

Further characterization of these tumour tissue compartments revealed that regions enriched for papillary and micropapillary growth patterns harboured amplifications on chromosome 1 (**Methods**), including the oncogene MYCL, linked with increased proliferation and metabolic activity, and the epigenetic regulator KDM4A, associated with enhanced invasiveness^37,38^ (**Figure 4c-d, Supplementary Figure 5**). This amplified region in the p-arm encompasses several additional cell cycle regulators including CDC20, E2F2, and CDCA8, collectively reinforcing proliferative and invasive hallmarks of cancer^39–41^. SLC2A1 (GLUT1), a critical glucose transporter gene essential for cellular responses to hypoxia^42^ was also amplified (**Figure 4d**), consistent with previous studies linking GLUT1 expression to micropapillary histology and invasive behavior in LUAD^43^. Together, these observations suggest that genomic alterations may contribute to shaping the transcriptional identity of the hypoxia-associated cell states in this patient’s tumour.

Similar to our observations in LUAD, spatial analysis of the BRCA Patient 2 tumour revealed two distinct regions with divergent histomorphology, each exhibiting localized enrichment of different cancer cell states (**Extended Data Figure 4a-c**). In this patient sample, one region was dominated by malignant cells with a focal amplification on chromosome 11q, including the oncogenic cell cycle regulator CCND1^44^, while the other region showed an arm-level gain of chromosome 8 (**Extended Data Figure 4d-g**).

In the BRCA Patient 3 biopsy, where clonal evolution was not evident based on CNV inference, the spatial distribution of cancer cell states was significantly correlated with local cancer cell density, suggesting that in the absence of substantial genomic divergence, microenvironmental factors may play a dominant role in shaping the emergence and localization of distinct malignant states (**Figure 4e-h, Supplementary Figure 6**).

In COAD, spatial mapping revealed cancer cell state localization associated with tumour invasion. In Patient 4, cancer cells associated with the *Translation-Active Proteostasis I* and *Translation-Active Stress* states were concentrated in more aggressive tumour regions closer to the muscularis externa layer of the colonic epithelium (**Figure 4i-k**). These cell states were further enriched in leading invasive fronts in two additional COAD samples, suggesting that translationally active stress-adaptive programs mark cancer cells occupying with local invasive tumour niches (**Extended Data Figure 5**).

Collectively, these spatial analyses reveal that transcriptionally defined cancer cell states exhibit non-random spatial organization linked to histological features, microenvironmental context, and underlying genomic alterations.

### Enrichment of Aggressive Cancer Cell States in Metastatic LUAD

As the leading cause of cancer-related mortality worldwide^45^, lung cancer underscores the urgent need to understand the mechanisms driving progression and metastasis. To study how cancer cell state composition changes during lung cancer progression, we applied ITHmapper to metastatic LUAD, associating each malignant cell to one of each 13 cell states previously defined in primary tumours (**Figure 2e**, **Figure 3b**). This allowed us to directly compare cell state proportions between metastatic and primary samples and assess whether specific states are selectively enriched following dissemination. Notably, four LUAD cell states were significantly enriched in metastatic samples (**Figure 5a**; Wilcoxon rank-sum test, P < 0.05), all previously linked to reduced patient survival (**Supplementary Table 4**). These included the *Proteasome-High Stress Responsive* and *Hypoxia Glycolytic EMT I* states, both of which were also spatially localized to micropapillary regions associated with invasiveness in LUAD Patient 1 (**Figure 4a,b**). Conversely, cell states such as *MHC-II Ciliated Metabolic*, which were not enriched in either primary or metastatic tumours, were significantly associated with longer survival (**Figure 5a, Supplementary Table 4**).

**Figure 5:**
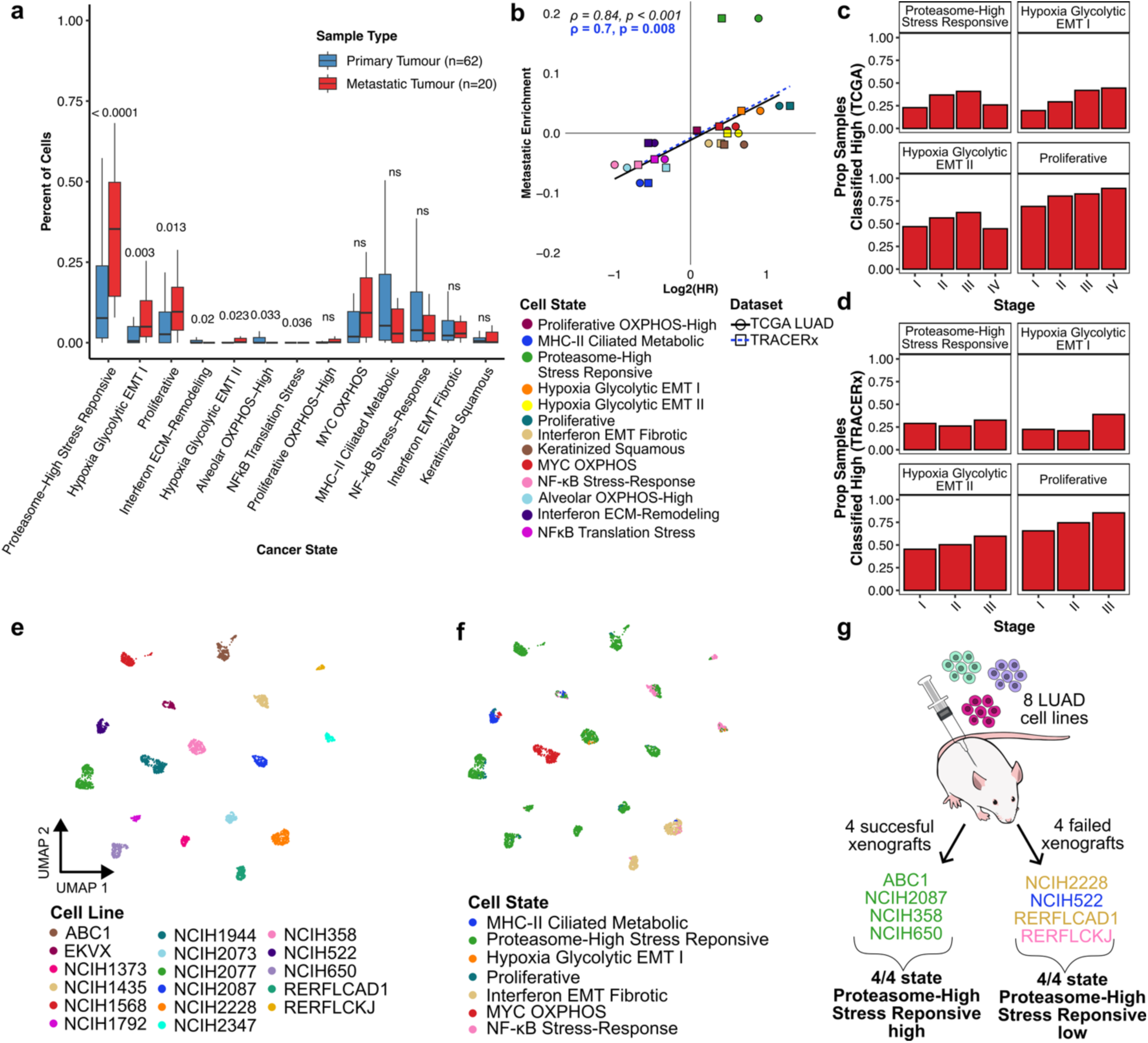
Enrichment of prognostic cell states in metastatic non-small cell lung adenocarcinoma. **a,** Comparison of the proportions of cell states among cancer cells in primary vs metastatic lung adenocarcinoma samples. Two-sided Wilcoxon rank sum test P values are indicated. Boxplot colours represent the sample type. Boxes and whiskers represent the lower fence, first quartile (Q1), median (Q2), third quartile (Q3), and upper fence. b, Correlation of poor-prognosis and metastatic enrichment in TCGA-LUAD (circles) and TRACERx-LUAD (squares). Metastatic enrichment was defined as the difference in the mean proportion of each cancer cell state between metastatic and primary samples. Points are coloured by mapped cell states. Spearman correlation coefficients from linear regression are shown on the plot. **c-d,** The proportion of samples (Prop) classified as high for various metastasis-enriched LUAD cell states across clinical pathological stages are shown for TCGA-LUAD (**c**) and TRACERx (**d**). Classifications were made based on the original optimal cut-point defined in the merged cohort containing all stages. **e-f,** UMAP visualization of lung adenocarcinoma cell lines derived from Kinker et al., 2020, coloured by cell line identity (**e**) and mapped cell state (**f**). **g,** Schematic representation of the comparison of xenograft success between cell lines exhibiting high versus low levels of the *Proteasome-High Stress Responsive* state. Only cell lines with corresponding scRNA-seq data (**e**) for cell state mapping are shown (Fisher exact test; P = 0.02857).

More broadly, we detected a strong positive correlation between the hazard ratios of LUAD cell states, derived from TCGA-LUAD survival data, and their scRNA-seq defined enrichment in metastatic versus primary tumours (Spearman ρ= 0.83, P < 0.001). This result was independently validated in the TRACERx-LUAD cohort (Spearman ρ= 0.7, P = 0.008) (**Figure 5b**).

Importantly, these associations were not solely attributable to clinical stage. Metastasis-enriched cell states were observed in both early- and late-stage LUAD samples, indicating that these aggressive phenotypes emerge in a substantial proportion of patients during the early stages of disease (**Figure 5c-d**).

To provide experimental support for the link between metastasis-enriched cancer cell states and metastatic potential, we analyzed xenograft data from LUAD cell lines^46^. Tumour-initiating capacity was used as a surrogate for metastatic colonization, based on the premise that successful engraftment reflects key features of dissemination and outgrowth at distant sites. Separately, we mapped LUAD cell lines profiled by scRNA-seq^47^ to their representative cell states using ITHmapper (**Figure 5e–f**). Among the subset of cell lines that overlapped with those used in the xenograft experiments, all lines predominantly composed of the metastasis-enriched *Proteasome-High Stress Responsive* state successfully formed xenografts (4/4), whereas those dominated by cell states not significantly enriched in metastatic tumours failed to engraft (4/4) (**Figure 5g**).

Together, our results demonstrate that LUAD cell states associated with poor survival and metastasis progression also exhibit tumour-initiating potential in vivo, identifying them as particularly clinically relevant and high-priority targets.

### Transcriptional cell states reveal actionable therapeutic vulnerabilities

To assess how malignant cell states respond to targeted therapies and to explore implications for individualized treatment strategies, we interrogated the Tahoe-100M dataset^48^, a large-scale single-cell perturbation resource in which a diverse panel of cancer cell lines was exposed to several hundred compounds and subsequently profiled by scRNA-seq. For each cancer type, we mapped both drug-treated and vehicle-treated (DMSO) cells onto our reference cell states. Next, for each drug-cell state pair, we assessed changes in cell state composition between treated and control samples to identify drugs that selectively depleted or enriched specific phenotypes (**Supplementary Table 5**). In these analyses, we focused on LUAD, COAD, and pancreatic adenocarcinoma (PAAD), each represented by at least five different cell lines in the Tahoe-100M dataset.

In LUAD, the proteasome inhibitor Bortezomib exhibited high potency across cell lines and showed particularly pronounced effects on the aggressive *Proteasome-High Stress Responsive* state (log_2_(OR) = -1.43, P < 1x10^-6^) previously associated with metastasis, and poor prognosis (**Figure 6a-d**). Globally, Bortezomib treatment induced marked shifts in transcriptomic profiles (**Figure 6c**), often accompanied by increased representation of less aggressive cell states such as *MHC-II Ciliated Metabolic* (log_2_(OR) = 5.82, P < 1x10^-25^) and *NFκB Stress-Response* (log_2_(OR) = 2.47, P < 1x10^-9^) (**Figure 6d, Extended Data Figure 6**).

**Figure 6:**
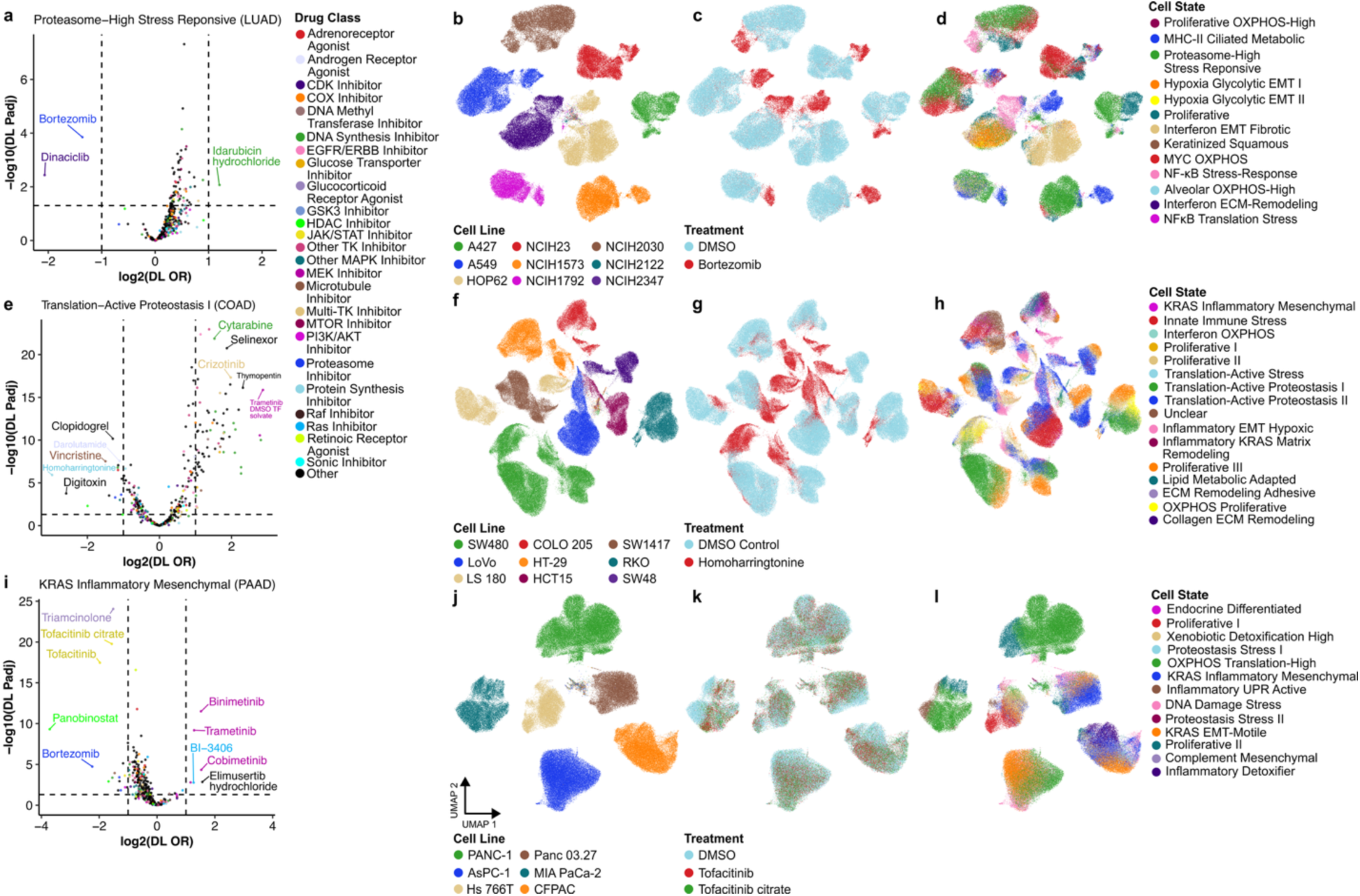
Therapeutic reprogramming of cell states across cancer types. **a,** Volcano plot showing pooled odds ratios for the *Proteasome-High Stress Responsive* state in LUAD under different drug treatments. **b–d,** UMAP visualizations of LUAD cell lines from the Tahoe-100M dataset, coloured by cell line identity (**b**), experimental treatment with Bortezomib or DMSO (**c**), and mapped cell state (**d**). **e**, Volcano plot showing pooled odds ratios for the *Translation-Active Proteostasis I* state in COAD under different drug treatments. **f–h,** UMAP visualizations of COAD cell lines, coloured by cell line identity (**f**), experimental treatment with Homoharringtonine or DMSO (**g**), and mapped cell state (**h**). **i**, Volcano plot showing pooled odds ratios for the *KRAS Inflammatory Mesenchymal* state in PAAD under different drug treatments. **j–l,** UMAP visualizations of PAAD cell lines, coloured by cell line identity (**j**), experimental treatment with Tofacitinib or DMSO (**k**), and mapped cell state (**l**). Pooled odds ratios were estimated using the DerSimonian-Laird random-effects model (DL-OR), reflecting the weighted combination of per-cell-line odds ratios for each drug within each cancer type. The top 10 significant drugs that preferentially enrich (log₂(DL-OR) > 1) or deplete (log₂(DL-OR) < –1) each cell state are labeled. Horizontal dashed lines indicate adjusted P < 0.05 (BH-corrected two-sided Wald Z-test). Drugs are coloured by their reported method of action in Tahoe-100M.

To further investigate therapeutic vulnerabilities of aggressive LUAD cell states, we analyzed PRISM drug repurposing data^49^, which measured viability of a broad panel of cancer cell lines following drug exposure. The analysis was focused on LUAD cell lines^47^ with high representation of the *Proteasome-High Stress Responsive* cancer state (**Figure 5e,f**). Among the 42 compounds showing selective toxicity toward these cell lines, candesartan, an angiotensin receptor blocker used to treat hypertension, showed the strongest selective activity (**Methods, Extended Data Figure 7**). This association highlights a potential opportunity to repurpose a well-tolerated drug to target a transcriptionally defined LUAD cell state associated with proteotoxic stress, metastatic capacity, and adverse clinical outcomes.

Collectively, these results nominate the proteasome as a promising therapeutic vulnerability in LUAD and suggest that patient stratification by cancer cell state composition, alongside rational combination strategies, may enhance the clinical efficacy of proteasome inhibitors, which have shown limited activity as monotherapies in prior trials^50,51^.

In COAD, treatment with the protein synthesis inhibitor, Homoharringtonine induced pronounced transcriptomic changes across cells and resulted in a significant depletion of the *Translation-Active Proteostasis I* cell state (log_2_(OR) = -2.98, P < 1x10^-7^) (**Figure 6e-h**), previously associated with invasive tumour fronts (**Figure 4 j-k, Extended Data Figure 5**).

In PAAD, treatment with the JAK/STAT inhibitors Tofacitinib and Tofacitinib Citrate, commonly used against inflammatory conditions such as arthritis and colitis, depleted inflammatory cancer cell states including *KRAS Inflammatory Mesenchymal* (Tofacitinib: log_2_(OR) = -1.98, P < 1x10^-19^, Tofacitinib Citrate: log_2_(OR) = -1.57, P < 1x10^-21^), and *Inflammatory Detoxifier* (Tofacitinib: log_2_(OR) = -1.12, P = 0.01, Tofacitinib Citrate: log_2_(OR) = -1.08, P < 1x10^-9^) (**Figure 6i-l**). Unlike the broader effects of Bortezomib and Homoharringtonine (**Figure 6c,g**), the global transcriptomic changes induced by Tofacitinib were more subtle, consistent with its targeted mechanisms of action involving specific pathway inhibition (**Figure 6k**).

### Cross-cancer therapeutic vulnerabilities

To identify broadly relevant therapeutic vulnerabilities recurring across multiple cancer types, we analyzed how drugs affect pan-cancer cell-state program defined by the pan cancer similarity network (**Figure 2f-g, Extended Data Figure 8**). Additionally, we grouped drugs based on their reported mechanisms of action and observed recurrent modulatory effects across mechanistically related compounds (**Extended Data Figure 9**). Notably, glucocorticoid agonists including Dexamethasone, Budenoside, and Bethamethasone-Dipropionate, significantly targeted the *Glycolytic Hypoxic Metabolism* program (**Extended Data Figure 8h**), typically identified in Bladder Urothelial Carcinoma (BLCA), BRCA, Glioblastoma Multiforme (GBM), High-Grade Serous Ovarian Carcinoma (HGSOC), Lower-Grade Glioma (LGG), and LUAD cancers (**Supplementary Table 2**).

A few drug classes, including HDAC and proteasome inhibitors, significantly targeted more than one pan-cancer program while simultaneously enriching others, highlighting their potential as versatile adjunctive agents with applicability across multiple cancer types. Notably, the *Neurotransmitter Release Cilia* program, shared among GBM, HGSOC, LGG, and STAD, was induced by several treatments, regardless of mechanism of action, but was depleted upon treatment with MEK inhibitors (**Extended Data Figure 9**). These findings underscore the need for additional combinatorial drug screens designed to identify and validate compounds capable of synergistically targeting all cancer cell states present within individual patients.

### Unsupervised clustering of drug-modulated transcriptional programs reveals potential mechanisms of action for uncharacterized compounds

To further explore drug effects across pan-cancer cell states, we performed hierarchical clustering of compounds based on their impact on distinct pan-cancer program types. This analysis revealed well-defined clusters of drugs with similar transcriptional modulation patterns (**Extended Data Figure 10**). Notably, compounds with unreported or ambiguous mechanisms of action clustered alongside well-characterized agents, suggesting related functional mechanisms. For instance, Clobetasol Propionate, a glucocorticoid receptor agonist drug^52^ annotated in Tahoe-100M as having an “unclear” mechanism, grouped with other established glucocorticoid receptor agonists Budesonide and Dexamethasone (**Extended Data Figure 10**). These results demonstrate that clustering of drug-induced transcriptional states provides a systematic framework to infer mechanisms of action for poorly characterized therapeutics, thereby informing future mechanistic studies and translational development.

## Discussion

The rapid expansion of scRNA-seq technologies has revolutionized our understanding of tumour heterogeneity across diverse cancer types. In this study, we assembled a pan-cancer atlas encompassing 3.6 million cells from over 1,000 tumours. Using a gene-gene covariance-based framework, we defined 259 discrete cellular states across 13 cancer types, representing 18 recurrent pan-cancer transcriptional programs. Examining both cancer type–specific and pan-cancer resolutions allowed us to uncover prognostic malignant cell states with previously unrecognized therapeutic vulnerabilities. Building on the cancer type–specific resolution, our approach integrates gene modules into discrete cell states that can be readily deconvoluted from bulk RNA-seq data, enhancing clinical translatability.

Unlike previous pan-cancer analyses^13,14^ that rely on expression matrix decomposition (e.g., non-negative matrix factorization), we identify gene modules based on covariance patterns in latent space, where transcriptomic proximity between cells is preserved. By leveraging gene-gene relationships within local cellular neighbourhoods of the phenotypic manifold, our approach reduces global variance, while enhancing the functional interpretability of the resulting transcriptional programs^7,22^. Although our framework emphasizes robust, generalizable cell states for clinical translation, we recognize it does not exhaustively capture rare subpopulations. The use of batch-corrected latent spaces mitigated technical effects between datasets but may have also removed more subtle patient specific biology, highlighting opportunities for further exploration.

Our analysis reaffirms the complexity and clinical significance of transcriptional diversity within tumours, while extending prior models of intratumoural heterogeneity^13,14^. Importantly, these transcriptional states are not abstract; when projected onto spatial transcriptomic and bulk RNA-seq data, they offer clinically useful insights. For instance, in LUAD, we identified hypoxia-driven and proteasome-activated cell states associated with poor survival, spatially localized to invasive regions, and enriched in metastatic samples^53–56^. In contrast, cell states linked to favorable prognosis were confined to acinar regions associated with moderate risk^57^. These data offer a molecular explanation for long-recognized pathological risk features.

Critically, we demonstrate that transcriptionally defined cancer cell states can provide prognostic resolution beyond conventional clinical criteria in established risk classification frameworks in NBL. This highlights the potential of single-cell transcriptomics to uncover clinically meaningful cellular programs whose signatures can be transferred to bulk transcriptomic data, facilitating the refinement of existing risk stratification systems. While this analysis centered on a single NBL-associated program, the *Proteostasis Stress* cell state, future studies will be needed to systematically assess the prognostic contribution of the cell states identified in this study across additional cancer types.

Spatial transcriptomic analyses conducted in this study further revealed that transcriptionally defined cancer cell states are not randomly distributed but instead align with distinct genetic features or architectural compartments within tumours. In the interrogated lung, breast, and colorectal cancers, malignant cell states exhibited spatial segregation, often demarcating histologically recognizable features such as papillary or acinar regions, tumour cores, or invasive fronts. These spatial patterns support the concept that transcriptional programs are tightly coupled to the local tumour microenvironment and underlying genomic alterations. Therapeutically, this spatial heterogeneity underscores the need to consider the regional context and composition of malignant phenotypes when designing targeted interventions.

To explore therapeutic vulnerabilities of aggressive LUAD states, we integrated PRISM drug screening data and high-throughput scRNA-seq from the Tahoe-100M dataset^48,49^. Proteasome inhibitors, most notably Bortezomib, selectively depleted the high-risk *Proteasome-High Stress Responsive* state while enriching less aggressive phenotypes. Although we cannot rule out clonal selection, we observed broad transcriptional shifts across all cell lines following Bortezomib treatment. Given that scRNA-seq in the Tahoe-100M dataset was performed just 24 hours post-treatment^48^, prior to substantial cell division, we interpret these changes as reflecting dynamic reprogramming rather than outgrowth of resistant clones. Future studies employing longitudinal single-cell transcriptomic profiling combined with lineage tracing and functional assays will be essential to dissect the temporal dynamics and reproducibility of drug-induced cell state transitions. Such approaches will enable accurate predictions of tumour transcriptomic trajectories and facilitate the rational design of drug combinations aimed at targeting emergent cell states.

In summary, our study presents a clinically oriented map of intratumoural heterogeneity, linking transcriptomic diversity to prognosis and treatment response. By connecting functionally distinct cell states with candidate therapeutics, we offer a blueprint for stratifying patients based on cell state composition and designing personalized treatment strategies tailored to specific malignant phenotypes. These insights move us closer to functional precision oncology, in which the transcriptional identity of cancer cells informs clinical decision-making and therapeutic development.

## Methods

### Pan-cancer single-cell atlas

#### Data curation

We collected publicly available human scRNA-seq datasets and metadata from published studies and repositories (Supplementary Table 1). Included studies met three criteria: (1) open access gene-expression count files, (2) cancer types covered by the scATOMIC training hierarchy^15^, and (3) scRNA-seq rather than single-nucleus RNA-seq (snRNA-seq). When available, sample-annotation tables were also retrieved to incorporate relevant clinical metadata.

#### Quality control

Raw gene-by-cell count matrices were filtered to remove cells expressing fewer than 500 genes or with >25% of reads mapping to mitochondrial genes. Gene names were standardized to Ensembl IDs using the ‘org.Hs.eg.db’ R package to ensure consistency across datasets. Doublets were predicted with the ‘scDblFinder’ R package^58^ using default parameters and removed following cell type annotation.

#### Cell type annotation

Cell type annotation was performed individually for each sample using scATOMIC v2.0.3^15^. Three parameter sets were applied: (1) confidence_cutoff = FALSE to maximize the number of cells receiving a final annotation, (2) use_CNVs = TRUE to perform CNV inference via the CopyKAT^16^ pipeline and identify high-confidence malignant cells, and (3) imputation = FALSE to minimize misclassifications potentially introduced by imputation. These complementary approaches were combined to define high-confidence malignant and non-malignant cells. Additionally, lower-resolution annotations were obtained by grouping related cells into their parental cell types within the scATOMIC hierarchy.

#### Data Integration and visualization

To handle the large dataset, count matrices were converted to on-disk format using the BPCells package^59^. Seurat-based sketching (RRID:SCR_016341)^60^ was applied for visualization purposes by sampling a representative subset of cells capturing the atlas diversity. Cells predicted as doublets or with doublet scores >0.5 were removed, and a Seurat object was created from the filtered counts. The object was split by cancer type and dataset, and up to 10,000 cells per layer were sketched using the ‘*SketchData’* function with the ‘*LeverageScore’* method. Data integration was performed using Seurat’s RPCA workflow, and the integrated embedding was projected onto the full dataset using ‘*ProjectIntegration’*. UMAP visualization was then applied to the integrated data. For cell subtype–specific integration, the same pipeline was used, but sketching was unnecessary due to smaller dataset sizes.

### Hotspot analysis

#### Cell and sample filtering

We interrogated the following cancer types: bladder urothelial carcinoma (BLCA), brain cancer (LGG/GBM), breast carcinoma (BRCA), colorectal adenocarcinoma (COAD/READ), gastric adenocarcinoma (STAD), kidney renal clear cell carcinoma (KIRC), liver hepatocellular carcinoma (LIHC), non-small cell lung adenocarcinoma (LUAD), neuroblastoma (NBL), ovarian high-grade serous carcinoma (HGSOC), pancreatic adenocarcinoma (PAAD), and prostate cancer (PRAD). The pan-cancer cell atlas was filtered to include only high-confidence malignant cells, defined by concordant scATOMIC malignant cell signature predictions and CopyKAT CNV inference. Additionally, cells annotated as blood or stromal cells when imputation = FALSE in scATOMIC were removed. Samples with fewer than 50 high-confidence malignant cells were excluded. Melanoma was omitted from all downstream analyses due to low number of primary tumour samples with >50 high-confidence malignant cells (n=4).

#### Data integration and batch effects correction

To mitigate technical batch effects, cancer type–specific datasets were integrated using a variational autoencoder implemented in scVI (RRID:SCR_026673)^23^. The scVI model was trained on the top 10,000 highly variable genes with flavor=“seurat_v3” parameter, using the sample identifier as the batch variable. Early stopping (early_stopping_patience = 10, early_stopping_min_delta = 10) was applied to prevent overfitting and ensure stable latent representations, while all other parameters were left at default. The latent space was extracted using ‘*get_latent_representation’*, and the integration was repeated 10 times with seeds 1–10 to ensure robustness.

#### Discovery of transcriptional programs in cancer cells

The Hotspot pipeline^22^ was applied separately to each cancer type, restricted to primary tumour samples. Data was renormalized in Scanpy (RRID:SCR_018139)^61^, and Hotspot was run on the top 2,000 highly variable genes using the ‘*danb*’ model and the scVI latent space described above. Autocorrelated features were calculated using the depth-adjusted negative binomial model, and a 30-nearest-neighbour graph was constructed. Genes with FDR < 0.05 were retained for local pairwise correlation calculations. For grouping genes into modules we set the following parameters: min_gene_threshold = 20, core_only = FALSE, and fdr_threshold = 0.01, as previous descibed^7^. Analyses were repeated 10 times with seeds 1–10 using the corresponding scVI replicates.

#### Quantifying module activity in individual cells

Hotspot module scores for each cell were computed using Hotspot’s implemented pipeline^22^. Briefly, a single-component PCA is performed on the expression of genes unique to each Hotspot module, and the resulting loadings are converted into scores, where negative values indicate low module activity and positive values indicate high activity. This procedure was repeated for all modules, and for each cell, the scores from the 10 replicate seeds mentioned above were combined into a single vector. To account for overlapping modules within and across seeds, Pearson correlations between module scores were clustered into meta-modules using the Ward D2 method. The resulting dendrogram was cut at the mean number of Hotspot modules across replicates. Singleton meta-module clusters, observed at only one replicate seed, which could represent random or non-reproducible programs, were excluded. The score for each meta-module in each cell was calculated as the mean of its constituent Hotspot modules.

#### Clustering meta-module scores

Cells were annotated with discrete cell states based on Louvain clustering of the nearest neighbour graph derived from meta-module scores at 10 different resolution parameters (0.1-1.0). Using multiple resolutions allowed the identification of cell states with varying granularity. To ensure clusters represented functionally coherent populations, cells with low silhouette scores across all resolutions were removed. Specifically, for each cell, silhouette scores were calculated using the ‘*ComputeSilhouetteScores’* function from the ‘SCISSORS’ R package^62^ and cells with a maximum silhouette score < 0.3 across all resolutions were excluded. Remaining cells were re-clustered at the 10 resolutions and clusters containing cells from fewer than 10% of samples within the corresponding cancer type were removed to eliminate highly patient-specific, non-generalizable states. The cells were re-clustered once more across all resolutions to assign a final label that was unaffected by the above filtered technical or patient specific effects.

#### Correlation of multi-resolution Hotspot clusters

To consolidate multi-resolution Louvain Hotspot-cluster labels into cell state annotations, clusters with correlated signals were merged. Specifically, we correlated the predicted fractions of cell states obtained from deconvolution of bulk RNA-seq datasets (TCGA/TARGET) across different clustering resolutions to identify distinguishable programs. Sarcoma was excluded from deconvolution and all downstream analyses due to a low number of patient samples in the bulk RNA-seq datasets (<100 samples in TARGET-Osteosarcoma). In brain and colorectal cancers, cell states were merged independently for GBM and LGG, and COAD and READ, respectively.

Each cancer-specific bulk dataset was deconvoluted using BayesPrism (RRID:SCR_027499)^24^ at each clustering resolution. For non-malignant cells, low-resolution scATOMIC annotations were used as cell types and high-resolution annotations were used as cell states. Low-level resolution cell types were defined as: cancer cells, B cells, dendritic cells, endothelial cells, fibroblasts, macrophages, mast cells, neutrophils, plasma cells, and T/NK cells. For non-malignant cells, terminal scATOMIC annotations were used as cell state. For the malignant cells the Hotspot-clusters were used as cell state. BayesPrism was run with ‘outlier.cut’ set to 0.01 and ‘outlier.fraction’ set to 0.1, without incorporating patient-specific information.

Pairwise Pearson correlations of predicted malignant Hotspot cluster proportions across the 10 clustering resolutions were computed. Louvain Hotspot-clusters with correlation values >0.8 were merged into meta-clusters, which were labeled according to the Louvain Hotspot-cluster containing the largest number of cells. Only meta-clusters encompassing more than three underlying cell states, representing robust, generalizable populations, were retained.

#### Intratumoural Transcriptomic Mapping of Heterogeneity Using ITHMapper

To assign each cell a final label corresponding to a single cancer cell state and to provide a reference framework for all downstream analyses as well as labeling cells in independent datasets, we calculated the Euclidean distance between each cell’s meta-module scores and the centroids of all meta-clusters. Centroids were defined as the mean score of each meta-module within the respective meta-cluster, and each cell was assigned to the cluster with the minimum distance.

To ensure a unified and reproducible framework for assigning query cells from independent datasets we applied this reference-based module-scoring and centroid-mapping strategy across all downstream analyses, including spatial localization, metastatic enrichment, drug perturbation, and xenograft studies. For each query cell, module scores were computed using the Hotspot modules defined in the primary pan-cancer single-cell atlas. The K-nearest neighbor graph was generated with 30 neighbours and here PCA was used as the latent space for spatial and cell line samples, which had lower batch complexity. Reference modules were appended to the Hotspot object rather than re-discovering new modules in the query dataset. Module scores were then calculated using Hotspot’s scoring pipeline, and each query cell was assigned to the meta-cluster centroid with the minimum Euclidean distance in module-score space.

#### Characterization of cell states using gene set enrichment analysis

Differential gene expression between cell states within each cancer type was assessed using a pseudobulk approach and was implemented in EdgeR (RRID:SCR_012802)^63^. Pseudobulk profiles were generated by summing gene expression counts for all cells within a given cell state for each sample. Two pseudobulk replicates per cell state per sample were created through random sampling of cells. Cell states with fewer than 15 cells in a sample were excluded. A ranked list of differentially expressed genes was generated using a quasi-likelihood fit EdgeR model with the cancer state and sample ID as design variables^63^. In LIHC and KIRC only cell state was used as a variable, as there were fewer samples. Gene set enrichment analysis was performed using the fgsea R package (RRID:SCR_020938)^64^ on a combined set of Hallmark (H) and Canonical Pathway (C2, CP, REACTOME, and KEGG_MEDICUS) gene sets from MSigDB (RRID:SCR_016863)^65^.

Each cell state was subsequently labeled based on its upregulated pathways, initially guided by a large language model and then manually refined.

#### Pan cancer similarity network

A pan-cancer similarity network was constructed with nodes representing individual cell states across cancer types. Edge weights were defined using a weighted Jaccard similarity based on pathway enrichment profiles. Specifically, each pathway in a cell state was assigned a weight equal to the negative logarithm of its adjusted p-value (−log(P adj.)). For each pair of cell states, the weighted Jaccard similarity was computed as follows: for every pathway, the minimum -log(P adj.) between the two states was summed to yield the intersection, and the maximum -log(P adj.) was summed to yield the union. The final edge weight was given by the ratio of the intersection to the union across all pathways for each pair (intersection/union). The network layout was visualized using the Fruchterman–Reingold force-directed algorithm implemented in the igraph R package (RRID:SCR_019225)^66^.

### Survival analysis

#### Deconvolution of bulk gene expression datasets

BayesPrism^24^ was used to deconvolute bulk RNA-seq and microarray datasets using cancer-specific cell states and normal cell types defined in the pan-cancer single-cell atlas. Data from 10,000 neutrophils^67^ were added to this reference as neutrophils are typically underrepresented in scRNA-seq studies. For non-malignant cells scATOMIC low-resolution annotations were used as cell types and high-resolution annotations were considered as cell states. Low-level resolution cell types were defined as: cancer cells, B cells, dendritic cells, endothelial cells, fibroblasts, macrophages, mast cells, neutrophils, plasma cells, and T/NK cells. For non-malignant cells, terminal scATOMIC annotations were used as cell state. For the malignant cells the meta-clusters were used as cell state. BayesPrism was run with ‘outlier.cut’ set to 0.01 and ‘outlier.fraction’ set to 0.1. Patient specific information was not provided to the BayesPrism model as we focused on recurrent cell states found across individuals. Whenever possible, raw count values were used. In samples where data was provided in log2, normalized format, the matrix was converted to unlogged format and normalized count values were used. Predicted cancer state proportions were normalized to the total percentage of malignant cell states, thereby correcting for variation in the proportion of non-malignant cells across samples.

#### Risk Stratification

For each cancer type, a discovery cohort was used to optimize the quantile split cut-point for stratifying patients into high- and low-risk groups. The cut-point was identified using the *surv_cutpoint* function from the survminer R package (RRID:SCR_021094)^68^, which selects the threshold that minimizes the log-rank P value. The minprop parameter was set to 0.25, ensuring that the smaller group contained at least 25% of patients. For adult cancers, the TCGA cohort served as the discovery set, while for neuroblastoma the TARGET cohort was used. Independent validation cohorts were then stratified using the same quantile threshold determined in the discovery cohort. All survival analyses were performed using 5-year overall survival as the outcome. Kaplan–Meier curves were generated with survminer^68^ and univariate hazard ratios were estimated using the *coxph* function in the survival R package^69^. P values were adjusted for multiple-testing using Benjamini-Hochberg correction within each cancer type and corresponding cohort used.

### Spatial Transcriptomic Analysis

Bin2Cell^36^ was used to assign multiple bins from the same cells into a single data point. Bins with zero expressed transcripts were discarded. Misalignment between the high-resolution microscope and Cytassist images was correct through manual alignment as described in the Bin2Cell documentation. Nuclei from high resolution microscopic images of the H & E slides were segmented using the built in ‘2D_versatile_he’ model included in ‘stardist’^70^. Fluorescent images from the CytAssist were segmented using the ‘2D_versatile_fluo’ model included in ‘stardist’^70^. Nuclei labels were expanded to a max bin distance of 4 in the lung and colorectal cancer samples.

In the breast cancer sample, the nuclei labels were expanded by a max bin distance of 1, given the higher density of nuclei. The corresponding cell by gene expression matrix was used for cell type annotation, cell state assignment, and CNV inference. Cell type annotation was performed using a combination of scATOMIC and manual annotation. Cells were clustered using the Leiden algorithm with resolution = 1. Count matrices were annotated using scATOMIC’s forced classification mode. Cells with sparse counts were observed to be randomly assigned scATOMIC labels with a bias towards T cells. To correct for this, the major cell type within a cluster was assigned to all cell types in the same respective cluster. Cluster cell type assignment was confirmed by comparing the top differentially expressed genes among each cluster. To differentiate cancer cells from normal tissue, visualization of the Leiden cluster on the respective histology was used. Clusters that were represented in regions of normal histology were annotated as “normal/unknown” cells. Subsequently, malignant cells were mapped to reference-defined cell states using ITHmapper. CNVs in Visium HD samples were inferred using ‘infercnvpy’ with the following parameters: ‘window_size’ = 250, ‘n_jobs’ = 20, ‘chunksize’ = 200, and calculate_gene_values = ‘True’. All non-malignant epithlelial, immune, and stromal cells were used as reference cell types. The corresponding result was filtered to only include malignant cells and CNV scores were subsequently clustered using the Leiden algorithm at resolution = 0.2 and 0.1 for the LUAD and BRCA samples respectively to identify potential subclones. To determine cancer cell density within regions of the breast cancer sample, the ‘calc.cell_density’ method from the Monkeybread^71^ Python module was used with approx = ‘True’ and resolution = ‘1’. Only cell states with greater than 10,000 cells in the sample were used in the cell state – cancer cell density association analysis.

### Metastatic enrichment analysis

Query malignant cells from both primary and metastatic malignant cell subsets of the LUAD data were mapped to primary-only reference-defined cell states using ITHmapper. Metastatic enrichment was defined as the difference in the mean proportion of each cancer cell state between metastatic and primary samples.

### Cell line xenograft and drug sensitivity analysis

Query cells were mapped to the reference-defined cell states using ITHmapper. Cell lines were considered high for the respective cell state that represented the majority of their cells. Eight cell lines from Kinker et al^47^ had corresponding xenograft success data in Matsubara et al^46^. Drug response data was retrieved from DepMap’s 24Q2 release of the PRISM screen (RRID:SCR_017655)^49^. Sensitivity was defined as the negative of the provided log2FC values. The comparison between cell lines high vs low in state *Proteasome-High Stress Responsive* was performed using the online Data Explorer tool version 2. Reported P values were calculated using an empirical-Bayes moderated t-test. Cell lines included in the high group were: ABC1 (RRID:CVCL_1066), EKVX (RRID:CVCL_1195), NCIH1373 (RRID:CVCL_1465), NCIH1435 (RRID:CVCL_1470), NCIH1568 (RRID:CVCL_1476), NCIH1792 (RRID:CVCL_1495), NCIH2073 (RRID:CVCL_1521), NCIH2087 (RRID:CVCL_1524), NCIH358 (RRID:CVCL_1559), NCIH650 (RRID:CVCL_1575). Cell lines included in the low group were: NCIH1944 (RRID:CVCL_1508), NCIH2228 (RRID:CVCL_1543), NCIH2347 (RRID:CVCL_1550), NCIH522 (RRID:CVCL_1567), RERFLCAD1 (RRID:CVCL_1651), RERFLCKJ (RRID:CVCL_1654). Data for the NCIH2077 (RRID:CVCL_5157) cell line was previously removed from DepMap due to inconsistent data fingerprints flagged in the 21Q3 release notes. Drugs were considered selectivity candidates if they had minimal effects on the low population (log2FC > -0.5) and inhibited growth in the high population (log2FC < -0.5). Additionally, the change in sensitivity between the two categories must be significant to categorize a drug as selective (Wilcoxon rank sum test, P<0.05).

### Drug-induced cell state dynamics using Tahoe-100M

To quantify the effect of drug treatment on LUAD cell states, query cells were mapped to reference-defined cell states using ITHmapper. For each drug and plate condition, we compared the abundance of each cell state between drug-treated and control DMSO wells. For each cell state in each cell line, we calculated an odds ratio and standard error using the Fisher test estimate as follows:

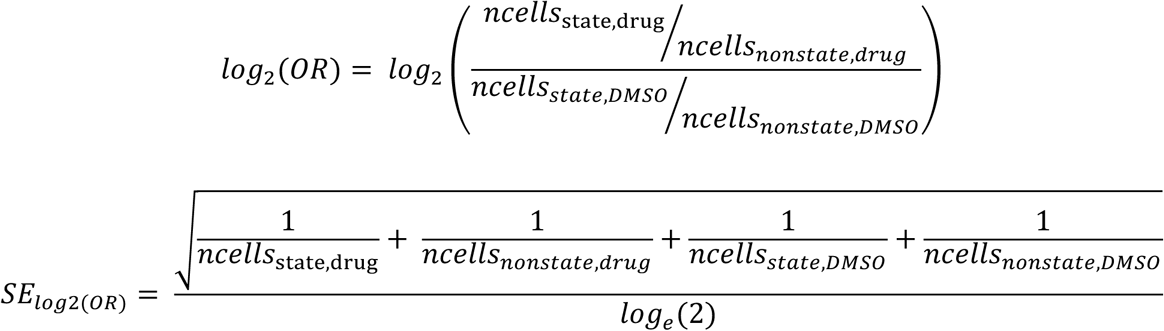

In cases where zeroes were present for any of the *ncells* values, to avoid infinite value and capture high or low odds ratios, we applied the Haldane correction, adding a value of 0.5 to each value.

To summarize drug effects across cell lines, we combined the log_2_OR values using the DerSimonian-Laird estimator via the rma function from the R ‘metafor’ package. Mechanism of action annotations were derived from metadata provided by the authors of Tahoe-100M.

## Data Availability

For convenience, we deposited the processed pan cancer single cell atlas along with its cell-level metadata in Zenodo^18^.

Accession numbers for the original, unprocessed datasets are described below. Single-cell datasets used for the pan cancer single cell atlas were originally retrieved from the following GEO deposited datasets: GSE115978, GSE131907, GSE132465, GSE134520, GSE137804, GSE140819, GSE141445, GSE148673, GSE151530, GSE152048, GSE154778, GSE161529, GSE165897, GSE176078, GSE178341, GSE183904, GSE185344.

Single-cell datasets used for the PCSCA were originally retrieved from the following Single Cell Portal deposited datasets: SCP1288, SCP1415, SCP2702.

Additional scRNA-seq datasets were downloaded directly from external sources described in the manuscripts: https://codeocean.com/capsule/8321305/tree/v1, https://datahub-262-c54.p.genap.ca/GBM_paper_data/GBM_cellranger_matrix.tar.gz, https://dna-discovery.stanford.edu/download/1403/, https://dna-discovery.stanford.edu/publicmaterial/datasets/mCRC_scRNAseq/mCRC_scRNA_filtered.zip, https://lambrechtslab.sites.vib.be/en/high-grade-serous-tubo-ovarian-cancer-refined-single-cell-rna-sequencing-specific-cell-subtypes, https://lambrechtslab.sites.vib.be/en/pan-cancer-blueprint-tumour-microenvironment-0, https://lambrechtslab.sites.vib.be/en/single-cell, https://static-content.springer.com/esm/art%3A10.1038%2Fs41467-020-18916-5/MediaObjects/41467_2020_18916_MOESM2_ESM.zip, https://www.nature.com/articles/s41467-022-29366-6#MOESM6, https://www.science.org/doi/suppl/10.1126/science.aat1699/suppl_file/aat1699_datas1.gz.zip, https://zenodo.org/record/3969339.

Additional scRNA-seq datasets were downloaded from HTAN via Synapse: https://data.humantumoratlas.org/publications/hta8_2021_cancer-cell_joseph-m-chan, HTAN BU, HTAN Vanderbilt, HTAN WUST, HTAPP, Moorman et al^7^ https://humantumoratlas.org/publications/hta8_2024_nature_a-r-moorman. scRNA-seq from Vazquez-Garcia et al^72^ was obtained from Synapse syn25569736.

Bulk gene expression datasets from TCGA^25^ and TARGET^26^ were downloaded from UCSC Xena (htseq_counts.tsv.gz). Bulk gene expression datasets from CPTAC were downloaded from https://proteomic.datacommons.cancer.gov/pdc/cptac-pancancer (RNA_Broad_V1.zip). Additional bulk gene expression datasets from were retrieved from the following GEO deposited datasets: GSE14520, GSE14764, GSE209746, GSE23554, GSE26712, GSE30161, GSE32894, GSE39582, GSE49711, or downloaded directly from external sources described in manuscripts: http://research-pub.gene.com/IMvigor210CoreBiologies/, https://data.mendeley.com/datasets/yzxtxn4nmd/4, https://static-content.springer.com/esm/art%3A10.1038%2Fs41591-020-0839-y/MediaObjects/41591_2020_839_MOESM2_ESM.xlsx, https://www.cbioportal.org/study/summary?id=coad_silu_2022, https://www.cbioportal.org/study/summary?id=paad_qcmg_uq_2016, https://zenodo.org/records/7603386.

Bulk gene expression the GLASS cohort was downloaded from Synapse syn17038081.

Visium HD data and images were directly downloaded from 10X Genomics’ datasets pages^32–35^: https://www.10xgenomics.com/datasets/visium-hd-cytassist-gene-expression-human-lung-cancer-post-xenium-expt, https://www.10xgenomics.com/products/visium-hd-spatial-gene-expression/dataset-human-crc, https://www.10xgenomics.com/datasets/visium-hd-cytassist-gene-expression-human-breast-cancer-fixed-frozen, https://www.10xgenomics.com/datasets/visium-hd-cytassist-gene-expression-libraries-human-breast-cancer-ff-ultima.

Cell line scRNA-seq data was downloaded from Single Cell Portal SCP542. Cell line scRNA-seq data from Tahoe-100M was downloaded as described in https://github.com/ArcInstitute/arc-virtual-cell-atlas/blob/main/tahoe-100M/tutorial-py.ipynb. Cell line xenograft data was retrieved from Table S3 in Matsubara et al^46^.

## Code Availability

The ITHmapper python module, associated code, and user manual are available at GitHub (https://github.com/abelson-lab/ITHmapper, https://github.com/abelson-lab/ITHmapper-manuscript) and deposited in Zenodo^18^ (https://doi.org/10.5281/zenodo.16852297).

## Description of Supplementary Data

**Supplementary Table 1: Overview of datasets employed for the pan-cancer single-cell atlas, bulk transcriptomics, and spatial transcriptomics**

**Supplementary Table 2: Gene set enrichment analysis and corresponding labels of malignant cell states across the pan cancer single-cell atlas.** Significantly upregulated gene sets from hallmark, KEGG_MEDICUS, and REACTOME pathways from MSigDB^65^ are shown for each meta-cluster. Manually curated cell state labels and program types are shown.

**Supplementary Table 3: Predicted cancer cell state fractions from deconvolution of bulk datasets.** Bars represent the predicted proportion of each cell state in each sample.

**Supplementary Table 4: Hazard ratios and corresponding P values for each cell state identified in the pan cancer single-cell atlas in cancer types with available bulk RNA seqeuncing and match clinical data.** Hazard ratios are given for the discovery datasets (TCGA/TARGET) and for validation datasets. Red and blue shading represent significant hazard ratios > 1 or < 1 respectively.

**Supplementary Table 5: Associations between drug treatment and cell states in the Tahoe-100M dataset.** Pooled odds ratios were estimated using the DerSimonian-Laird random-effects model, reflecting the weighted combination of per-cell-line odds ratios for each drug within each cancer type.

## Acknowledgements

This work was supported by Investigator Awards received from the Ontario Institute for Cancer Research with funds from the province of Ontario to S.A. and P.A. I.N.M. obtained funds from the Ontario Graduate Scholarship Program – University of Toronto.

## Author Information

All authors discussed the results, wrote, and commented on the paper. I.N.M. developed the concept, performed data analysis, and wrote the code. A.C. contributed to spatial CNV calling.

T.W.O. contributed to the pan cancer similarity network and intellectual discussions. S.H.B. provided pathological annotations for the spatial transcriptomics samples. S.A. and P.A. conceived the idea, contributed to data analysis, led, and supervised all aspects of the study.

## Ethics Declarations

The authors declare no competing interests.

---

**Extended Data Figure 1:**
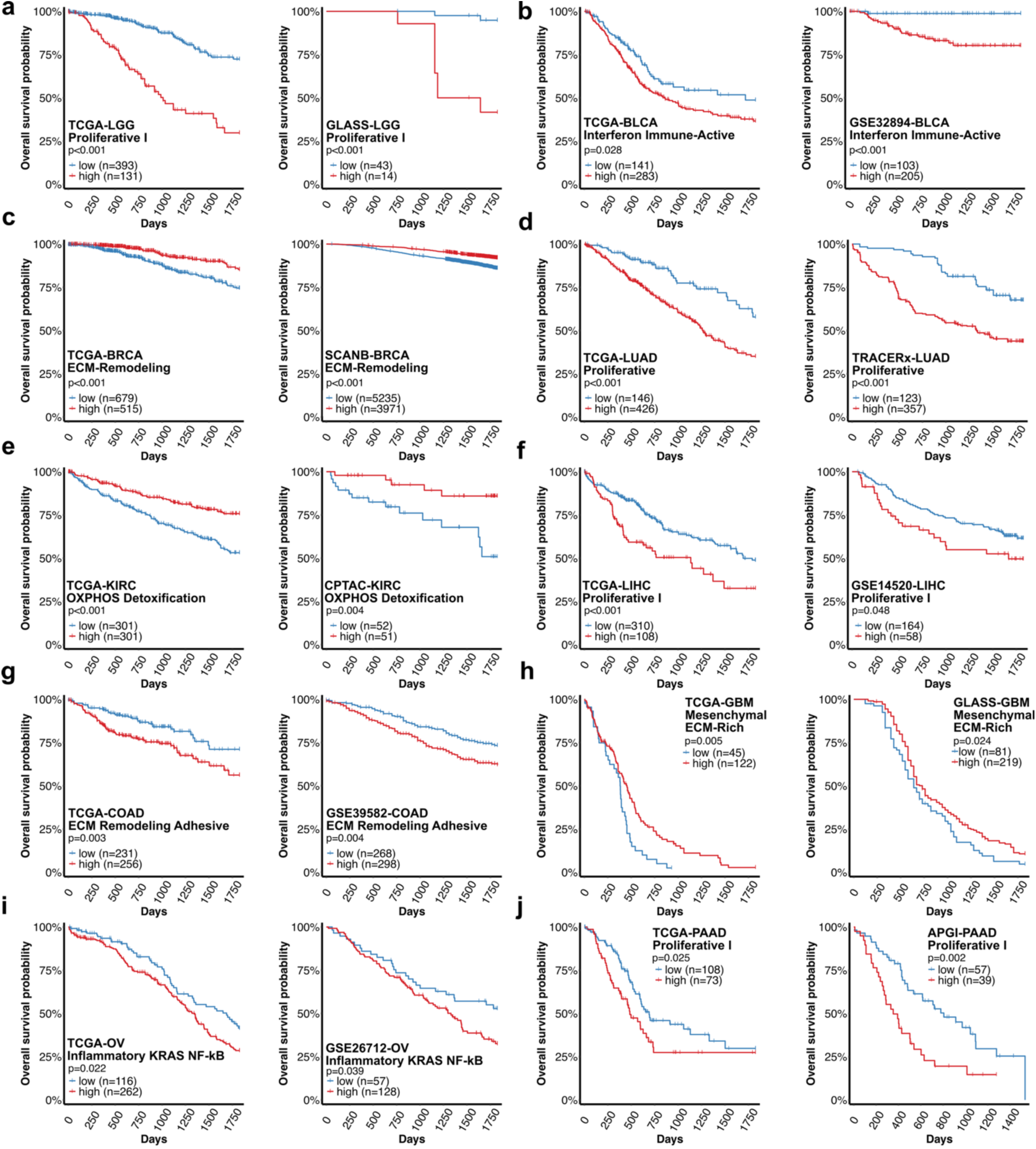
Cell state deconvolution predicts 5-year survival across cancer types. **a-j,** Kaplan-Meier plots for the top prognostic cell state in each cancer type: Low-Grade Glioma (LGG, **a**), Bladder Urothelial Carcinoma (BLCA, **b**), Breast Invasive Carcinoma (BRCA, **c**), Lung Adenocarcinoma (LUAD, **d**), Kidney Renal Clear Cell Carcinoma (KIRC, **e**), Liver Hepatocellular Carcinoma (LIHC, **f**), Colon Adenocarcinoma (COAD, **g**), Glioblastoma Multiforme (GBM, **h**), Ovarian Serous Cystadenocarcinoma (OV, **i**), and Pancreatic Adenocarcinoma (PAAD, **j**), shown for both TCGA/TARGET and a second validation cohort. All analyses plotted were significant by log-rank test (P < 0.05) in both cohorts. Red lines indicate the cell-state high group, and blue lines indicate the cell-state low group. Sample sizes for each group are indicated.

**Extended Data Figure 2:**
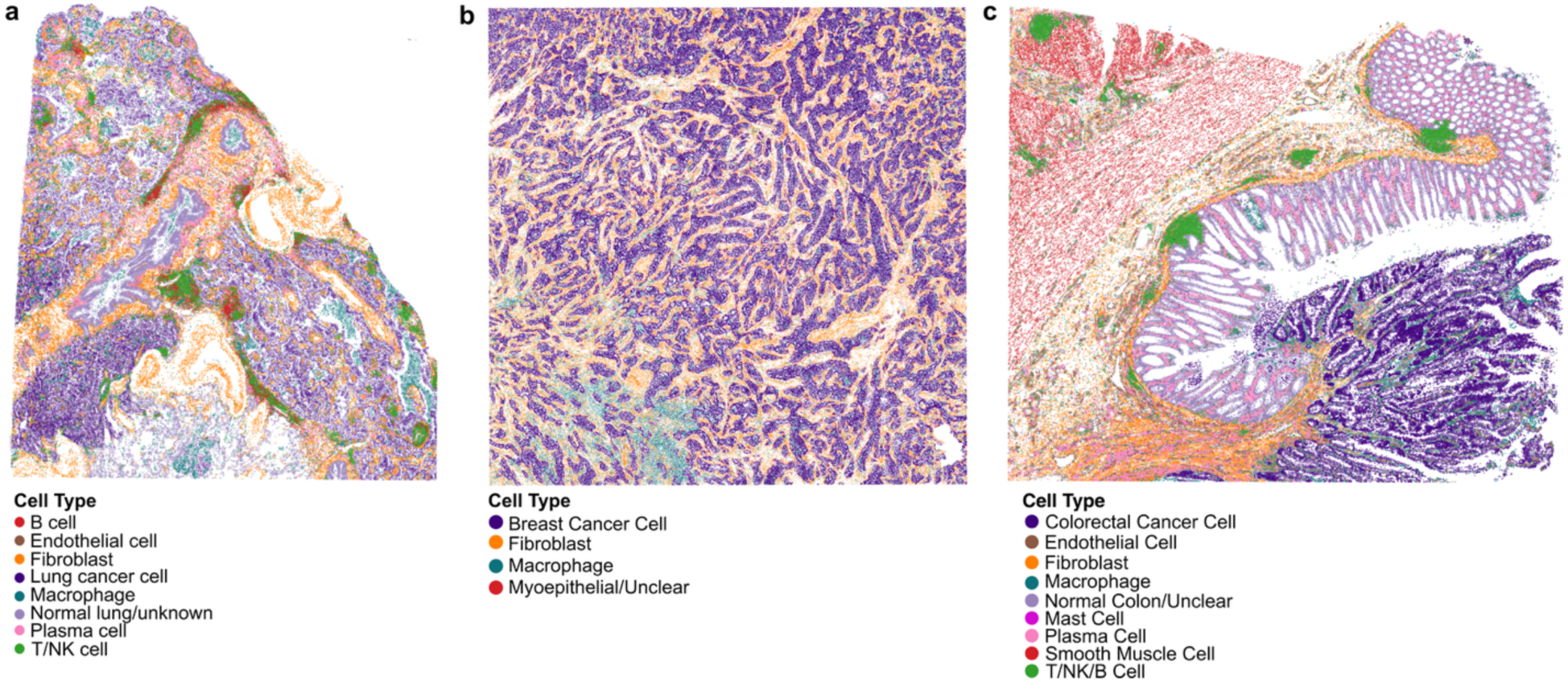
Spatial maps of predicted cell types in Visium HD samples. Cells are coloured by predicted cell type in LUAD **(a)**, BRCA **(b)**, and COAD **(c)**. Cell type assignments were derived from scATOMIC predictions, refined by manual marker-based annotation and histological assessment.

**Extended Data Figure 3:**
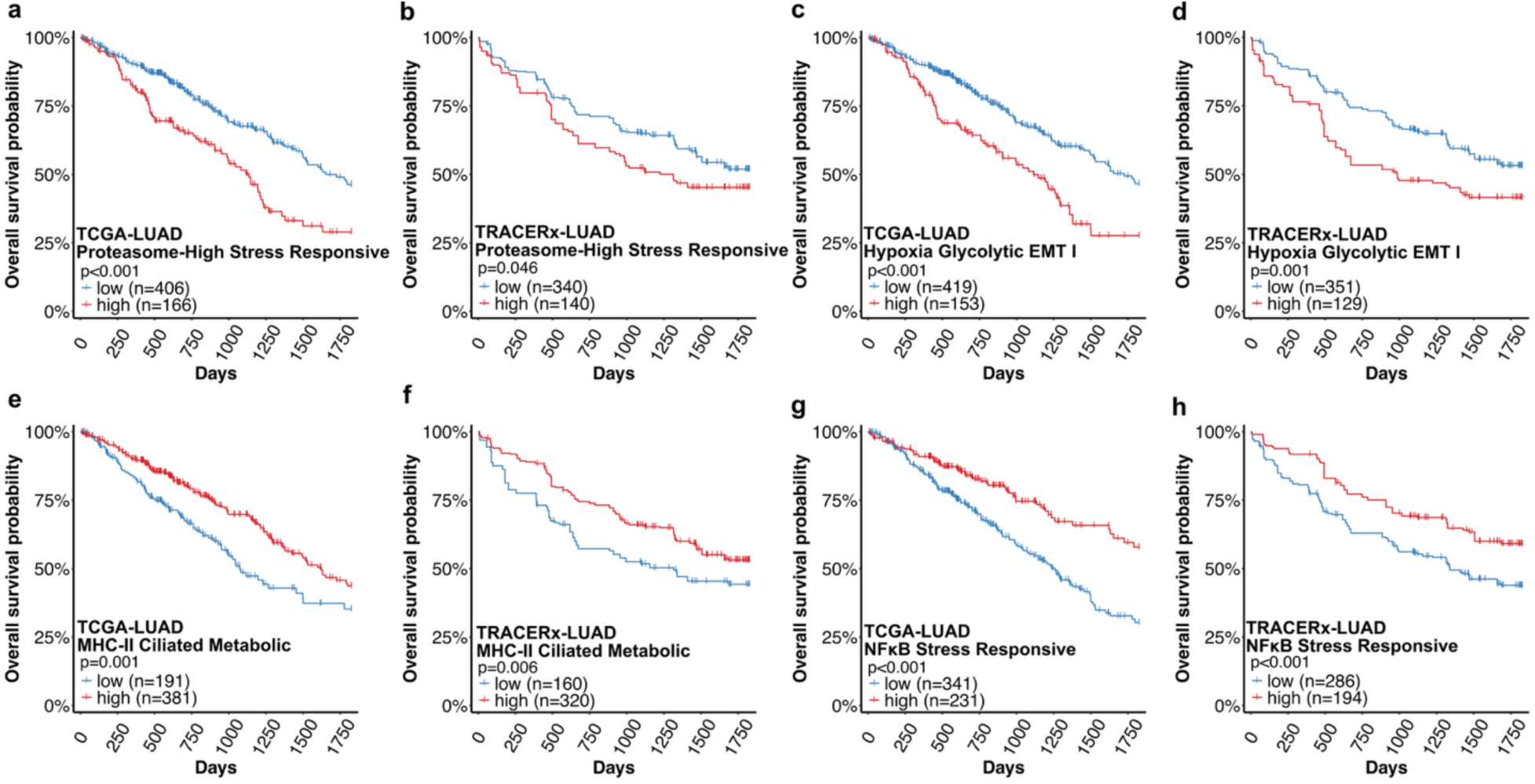
Survival analyses of spatially localized LUAD cell states. Kaplan-Meier plots for spatially localized LUAD cell states in the TCGA-LUAD and TRACERx-LUAD cohorts. Analyses shown had log-rank test P < 0.05 in both cohorts. Patients were stratified into cell-state high and low groups based on inferred abundance of each cancer cell state. The upper four panels (**a–d**) show cell states associated with inferior outcomes, while the bottom four panels **(e–h**) show cell states associated with favorable prognosis.

**Extended Data Figure 4:**
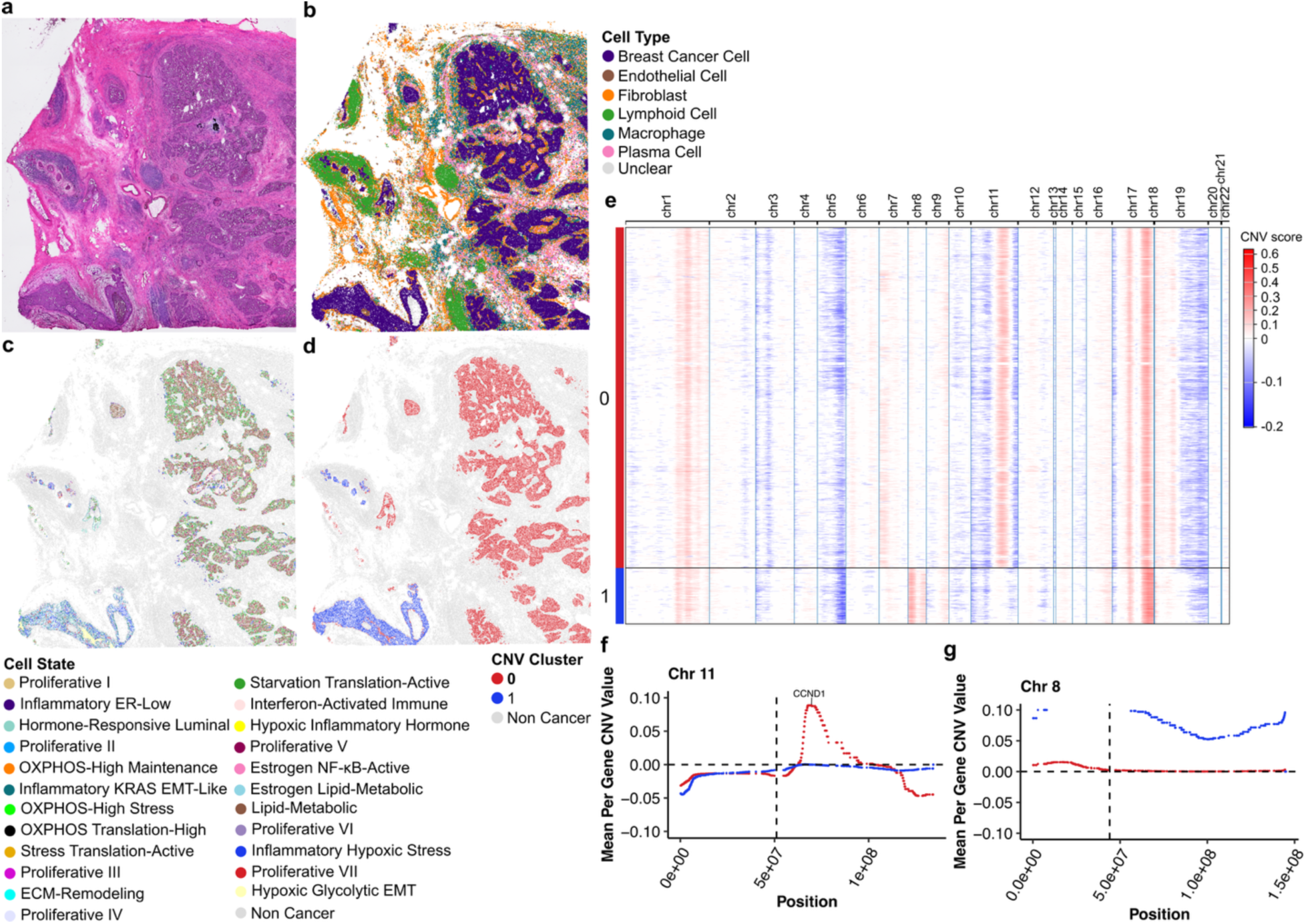
Spatial localization of cell states and copy number alterations in Visium HD breast cancer sample from Patient 2. **a,** Hematoxylin and eosin (H&E) staining of breast cancer Patient 2. **b-d,** spatial maps of Visium HD cells coloured by cell type (**b**), mapped cancer cell state (**c**), and CNV cluster (**d**). **e,** Heatmap of CNV scores in malignant cells with red and blue indicating gains and losses respectively. Leiden clusters of CNV profiles are shown on the y-axis. **f-g,** Mean per-gene CNV values along chromosome 11 (**f**) and chromosome 8 (**g**) for each CNV cluster (0 or 1).

**Extended Data Figure 5:**
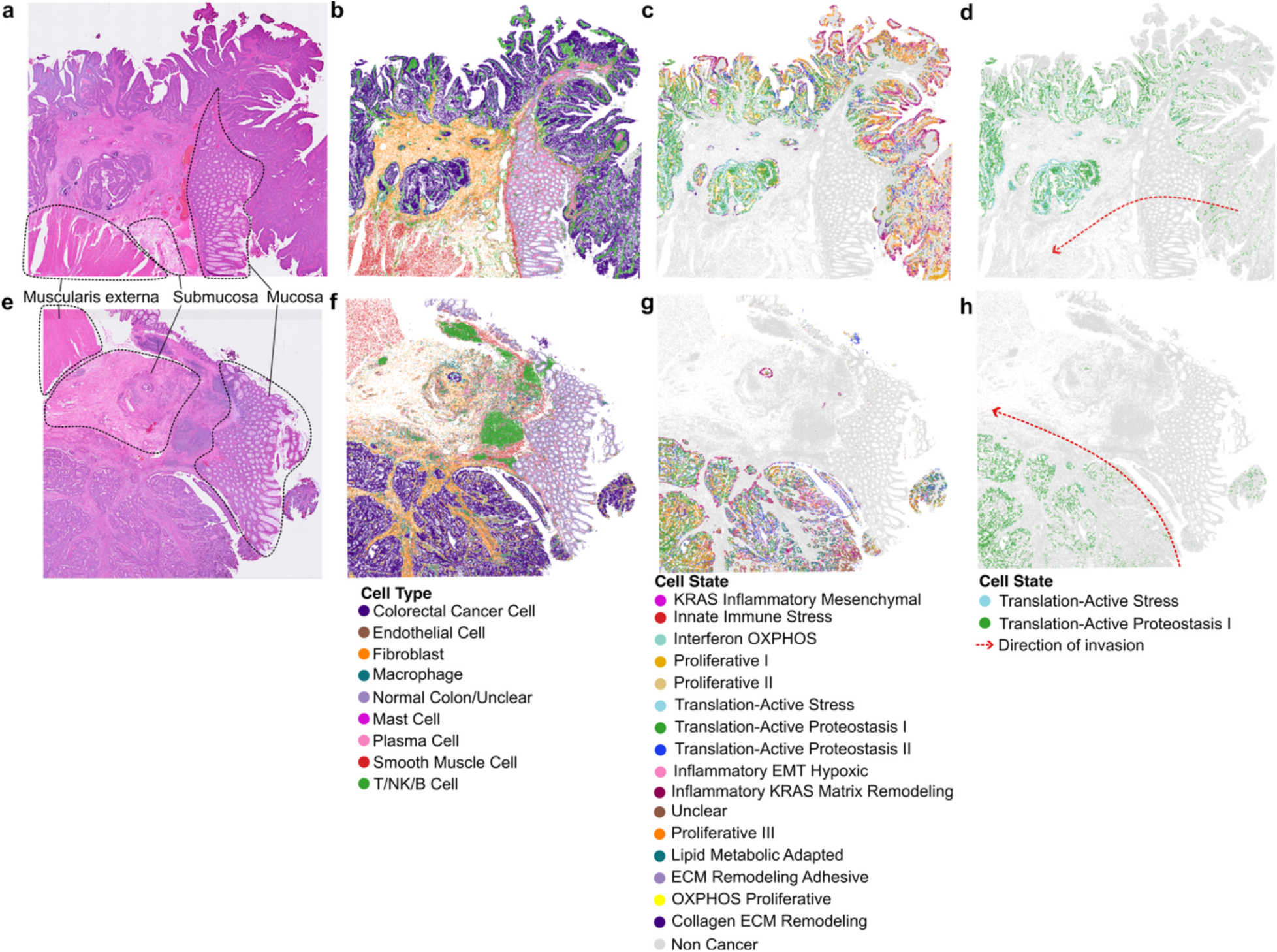
Spatial localization patterns of cancer cell states in colorectal carcinoma Visium HD samples. **a,** Hematoxylin and eosin (H&E) staining of COAD Patient 5. **b-c,** Spatial maps of cells from COAD Patient 5 coloured by cell type (**b**) and cell state (**c**). **d,** H&E staining of COAD Patient 6. **e-f,** Spatial maps of cells from COAD Patient 6 coloured by cell type (**e**) and cell state (**f**).

**Extended Data Figure 6:**
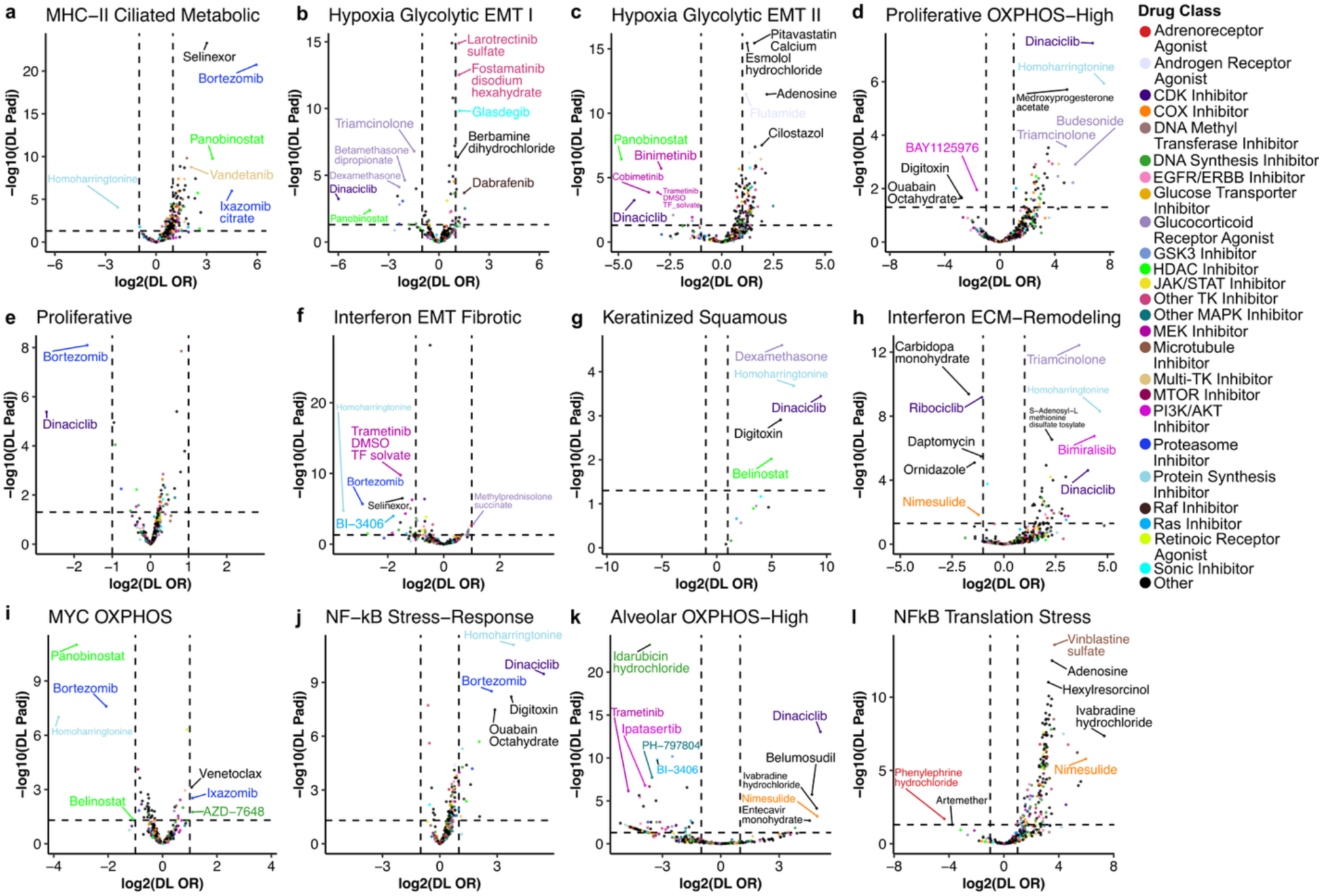
Volcano plots of odds ratio across cell lines for additional LUAD cell states. Volcano plots showing DerSimonian-Laird-estimated pooled odds ratio of cell state proportions under different drug treatments. Pooled odds ratios represent the weighted combination of per-cell-line odds ratios for each drug. The top 5 significant cell-state enriching (log_2_(DL OR) > 1) and depleting (log_2_(DL OR) < -1) drugs are labeled. Horizontal dashed lines indicate adjusted P = 0.05 (BH-corrected two-sided Wald Z-test). Drugs are coloured by their reported method of action in Tahoe-100M. Figure panels represent different LUAD cell states.

**Extended Data Figure 7:**
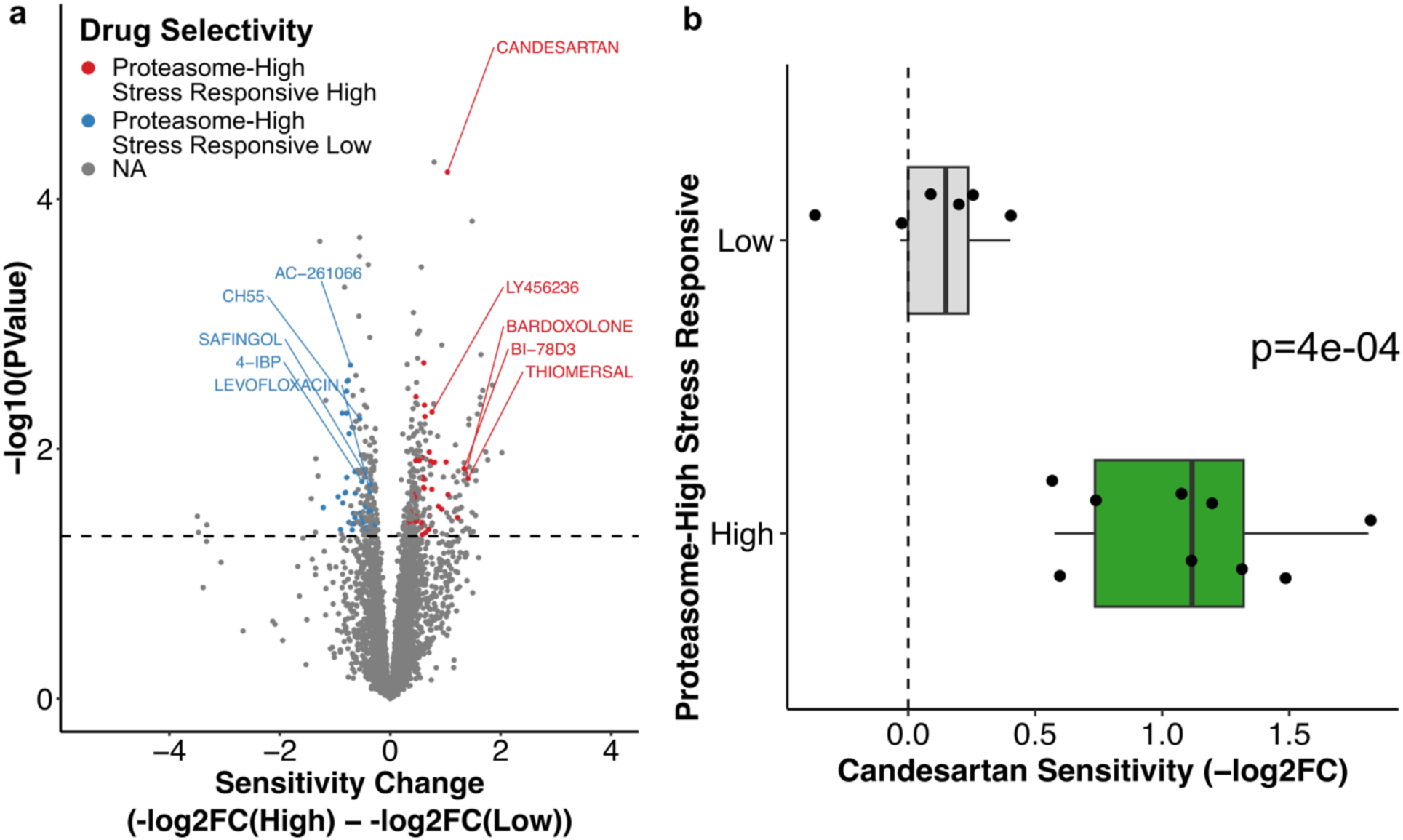
Aggressive cell states are selectively susceptible to repurposed drugs. **a,** Volcano plot comparing drug sensitivity in LUAD cell lines with high versus low proportions of the *Proteasome-High Stress Responsive* cell state. Positive effect sizes indicate increased drug susceptibility in cell lines classified as “high”. Red and blue dots denote drug candidates for selectively targeting high and low cell lines respectively. Drugs were considered selective candidates if they had minimal inhibitory effects on the cell state-low population (log_2_FC > -0.5) and inhibited growth in the cell state-high population (log_2_FC < -0.5) and vice versa. Drugs that did not meet the criteria for selectivity are labeled as NA (coloured gray). The top 5 candidates ranked by P value and effect size are labeled. The horizontal dashed line indicates adjusted P = 0.05 (BH-corrected empirical-Bayes moderated t-test, retrieved from the DepMap Explorer Tool). **b,** Comparison of the effect of candesartan on growth in cell lines high vs low for the Proteasome-High Stress Responsive cell state. Boxes and whiskers represent the lower fence, first quartile (Q1), median (Q2), third quartile (Q3), and upper fence. P value was calculated using a two-sided Wilcoxon rank-sum test.

**Extended Data Figure 8:**
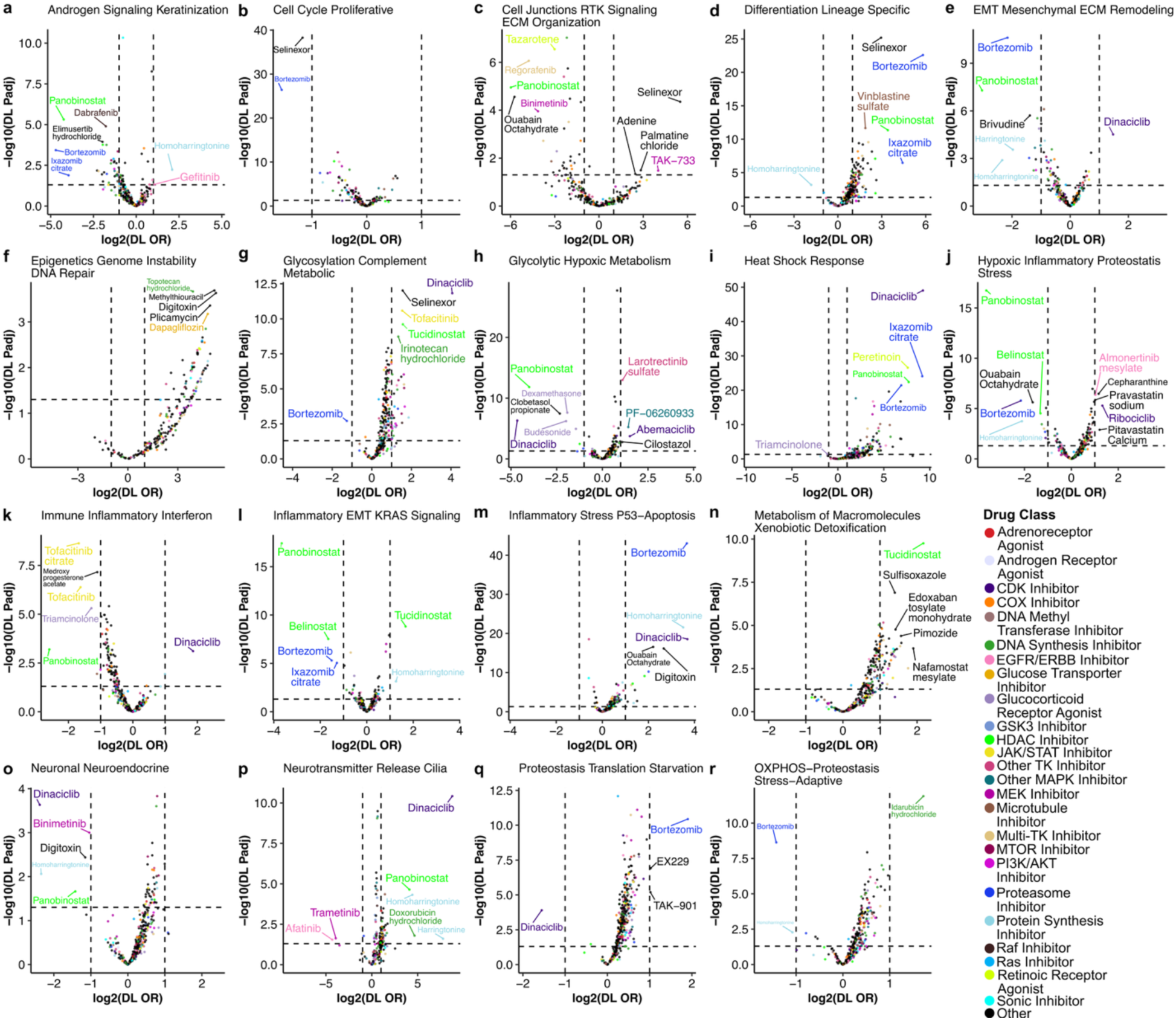
Volcano plots of odds ratio across pan cancer program types and drug classes. Volcano plots showing DerSimonian-Laird-estimated pooled odds ratios of pan cancer program proportions under different drug treatments. Pooled odds ratios represent a combination of per-cell-line odds ratios for each drug. Horizontal dashed lines indicate adjusted P = 0.05 (BH-corrected two-sided Wald Z-test). The top 5 significant cell state-enriching (log_2_(DL OR) > 1) and depleting (log_2_(DL OR) < -1) drugs are labeled. Drugs are coloured by their reported method of action in Tahoe-100M. Figure panels **a–r** represent different pan cancer cell state program types.

**Extended Data Figure 9:**
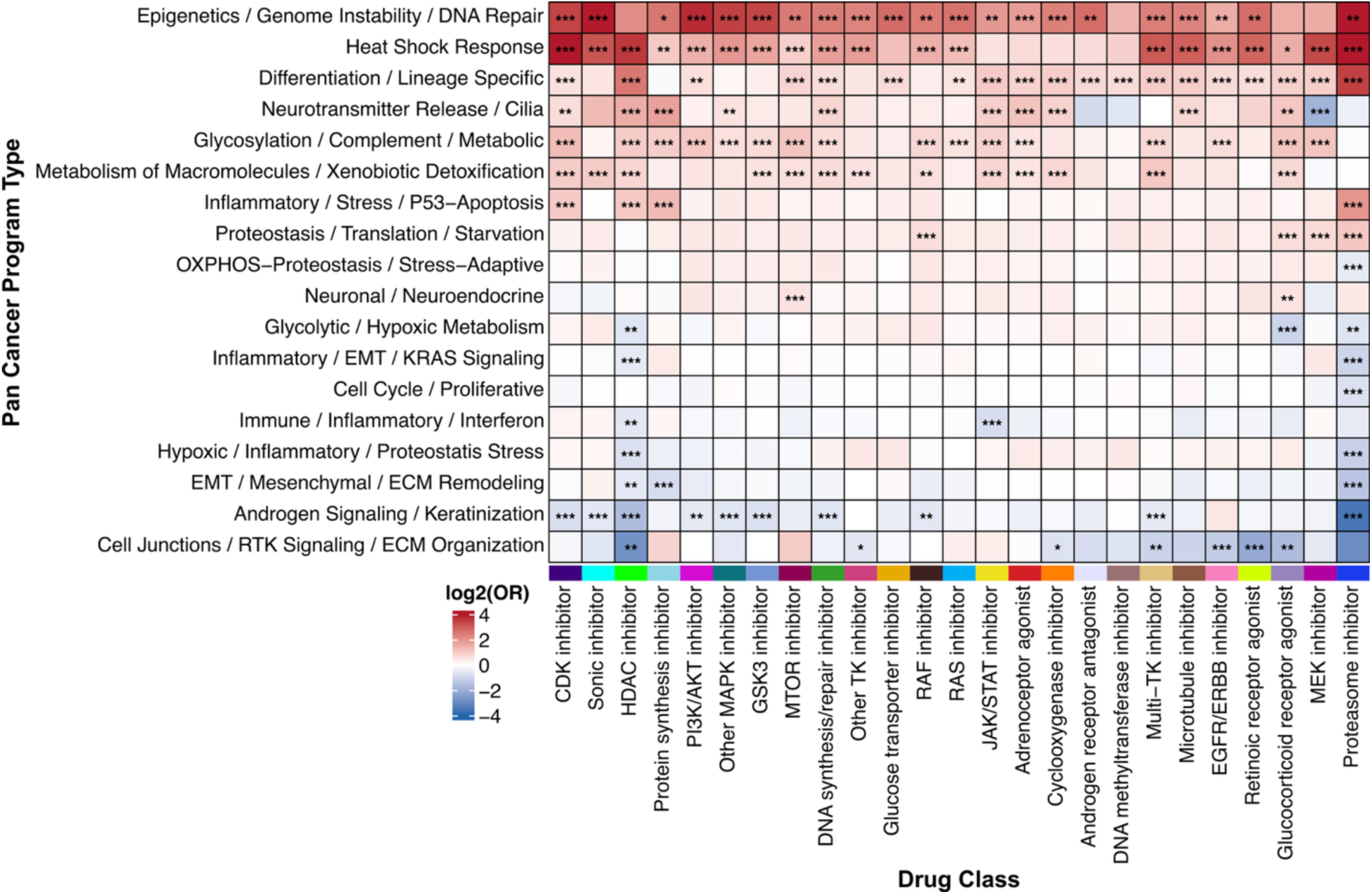
Recurrent modulatory effects of mechanistically related compounds on pan-cancer program types. Heatmap of DerSimonian-Laird-estimated pooled odds ratio of pan cancer program types in pan cancer cell lines treated in the presence of different drugs. Pooled odds ratios represent a combination of all treatments from drugs with the same reported method of action. Adjusted P values represent BH-corrected two-sided Wald Z-test, *** P < 0.001, ** P < 0.01, * P < 0.05. The heatmap tiles are coloured by the log_2_(DL ORs) with red and blue indicating drug classes that induce and deplete program types respectively.

**Extended Data Figure 10:**
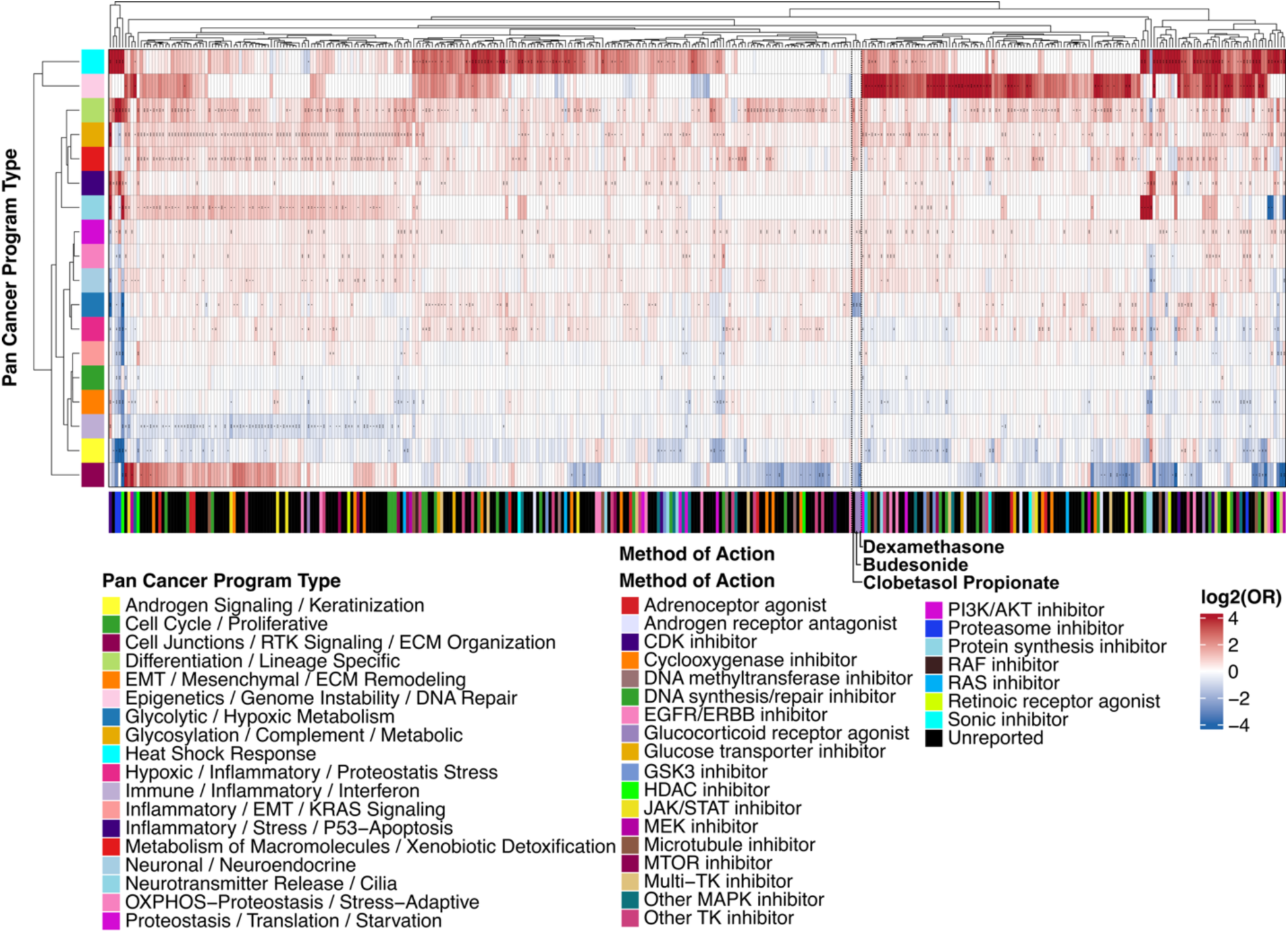
Modulatory effects of various compounds on pan-cancer program types. Heatmap showing DerSimonian-Laird-estimated pooled odds ratio (DL OR) for pan-cancer programs across treated cell lines. Pooled odds ratios represent a weighted combination of per-cell-line effects for each compound. Clobetasol Propionate, Budesonide, and Dexamethasone, an example of co-clustering drugs, are indicated on the plot with labels and a dotted box. Drugs with black labels indicate unreported mechanisms of action in Tahoe-100M metadata. Adjusted P values represent BH-corrected two-sided Wald Z-test, *** P < 0.001, ** P < 0.01, * P < 0.05. Heatmap tiles are coloured by the log_2_(DL ORs) with red and blue indicating drugs that induce and deplete program types respectively.

## References

1. Marusyk, A., Janiszewska, M. & Polyak, K. Intratumor Heterogeneity: The Rosetta Stone of Therapy Resistance. Cancer Cell 37, 471–484 (2020).

2. Tirosh, I. & Suva, M. L. Cancer cell states: Lessons from ten years of single-cell RNA-sequencing of human tumors. Cancer Cell 42, 1497–1506 (2024).

3. Baron, M. et al. The Stress-Like Cancer Cell State Is a Consistent Component of Tumorigenesis. cels 11, 536–546.e7 (2020).

4. França, G. S. et al. Cellular adaptation to cancer therapy along a resistance continuum. Nature 631, 876–883 (2024).

5. Puram, S. V. et al. Cellular states are coupled to genomic and viral heterogeneity in HPV-related oropharyngeal carcinoma. Nat Genet 55, 640–650 (2023).

6. Jerby-Arnon, L. et al. A Cancer Cell Program Promotes T Cell Exclusion and Resistance to Checkpoint Blockade. Cell 175, 984–997.e24 (2018).

7. Moorman, A. et al. Progressive plasticity during colorectal cancer metastasis. Nature 637, 947–954 (2025).

8. Valdes-Mora, F. et al. Single-cell transcriptomics in cancer immunobiology: The future of precision oncology. Frontiers in Immunology 9, (2018).

9. Neftel, C. et al. An Integrative Model of Cellular States, Plasticity, and Genetics for Glioblastoma. Cell 178, 835–849.e21 (2019).

10. Chan-Seng-Yue, M. et al. Transcription phenotypes of pancreatic cancer are driven by genomic events during tumor evolution. Nat Genet 52, 231–240 (2020).

11. Kim, N. et al. Single-cell RNA sequencing demonstrates the molecular and cellular reprogramming of metastatic lung adenocarcinoma. Nature Communications 2020 11:1 11, 1–15 (2020).

12. Wu, S. Z. et al. A single-cell and spatially resolved atlas of human breast cancers. Nature Genetics 2021 53:9 53, 1334–1347 (2021).

13. Barkley, D. et al. Cancer cell states recur across tumor types and form specific interactions with the tumor microenvironment. Nat Genet 54, 1192–1201 (2022).

14. Gavish, A. et al. Hallmarks of transcriptional intratumour heterogeneity across a thousand tumours. Nature 618, 598–606 (2023).

15. Nofech-Mozes, I., Soave, D., Awadalla, P. & Abelson, S. Pan-cancer classification of single cells in the tumour microenvironment. Nature Communications 2023 14:1 14, 1–14 (2023).

16. Gao, R. et al. Delineating copy number and clonal substructure in human tumors from single-cell transcriptomes. Nature Biotechnology 1–10 (2021) doi:10.1038/s41587-020-00795-2.

17. Virshup, I., Rybakov, S., Theis, F. J., Angerer, P. & Wolf, F. A. anndata: Access and store annotated data matrices. Journal of Open Source Software 9, 4371 (2024).

18. Nofech-Mozes, I. Data and codes for: A Pan-Cancer Single-Cell Compendium of Intratumoural Heterogeneity. 10.5281/zenodo.16852297 (2025) 10.5281/zenodo.16852297.

19. Langfelder, P. & Horvath, S. WGCNA: an R package for weighted correlation network analysis. BMC Bioinformatics 9, 559 (2008).

20. Aibar, S. et al. SCENIC: single-cell regulatory network inference and clustering. Nat Methods 14, 1083–1086 (2017).

21. Kotliar, D. et al. Identifying gene expression programs of cell-type identity and cellular activity with single-cell RNA-Seq. eLife 8, e43803 (2019).

22. DeTomaso, D. & Yosef, N. Hotspot identifies informative gene modules across modalities of single-cell genomics. Cell Syst 12, 446–456.e9 (2021).

23. Lopez, R., Regier, J., Cole, M. B., Jordan, M. I. & Yosef, N. Deep generative modeling for single-cell transcriptomics. Nat Methods 15, 1053–1058 (2018).

24. Chu, T., Wang, Z., Pe’er, D. & Danko, C. G. Cell type and gene expression deconvolution with BayesPrism enables Bayesian integrative analysis across bulk and single-cell RNA sequencing in oncology. Nat Cancer 3, 505–517 (2022).

25. Weinstein, J. N. et al. The Cancer Genome Atlas Pan-Cancer analysis project. Nat Genet 45, 1113–1120 (2013).

26. Pugh, T. J. et al. The genetic landscape of high-risk neuroblastoma. Nat Genet 45, 279–284 (2013).

27. Kolberg, L., Raudvere, U., Kuzmin, I., Vilo, J. & Peterson, H. gprofiler2 -- an R package for gene list functional enrichment analysis and namespace conversion toolset g:Profiler. F1000Res 9, ELIXIR-709 (2020).

28. Newman, A. M. et al. Determining cell type abundance and expression from bulk tissues with digital cytometry. Nature Biotechnology 37, 773–782 (2019).

29. Vermeulen, J. et al. Predicting Outcomes for Children with Neuroblastoma. Discovery Medicine 10, 29–36 (2010).

30. Liang, W. H. et al. Tailoring Therapy for Children With Neuroblastoma on the Basis of Risk Group Classification: Past, Present, and Future. JCO Clin Cancer Inform 895–905 (2020) doi:10.1200/CCI.20.00074.

31. Zhang, W. et al. Comparison of RNA-seq and microarray-based models for clinical endpoint prediction. Genome Biol 16, 133 (2015).

32. Oliveira, M. F. de et al. High-definition spatial transcriptomic profiling of immune cell populations in colorectal cancer. Nat Genet 57, 1512–1523 (2025).

33. 10X Genomics. Visium HD Spatial Gene Expression Libraries, Post-Xenium, Human Lung Cancer (FFPE), Experiment 2, HD only (control), HD Spatial Gene Expression dataset analyzed using Space Ranger 3.0.0. (2024).

34. 10X Genomics. Visium HD Spatial Gene Expression Library, Human Breast Cancer (Fixed Frozen), HD Spatial Gene Expression dataset analyzed using Space Ranger 3.1.1. (2024).

35. 10X Genomics. Visium HD Spatial Gene Expression Library, Human Breast Cancer (Fresh Frozen), Ultima Sequencing, HD Spatial Gene Expression dataset analyzed using Space Ranger 3.1.2. (2025).

36. Polański, K. et al. Bin2cell reconstructs cells from high resolution Visium HD data. Bioinformatics 40, btae546 (2024).

37. Jing, Q., Jun, X. F., Hui, L. C., Na, H. & Yang, L. H. MYCL1 Amplification and Expression of L-Myc and c-Myc in Surgically Resected Small-Cell Lung Carcinoma. Pathol. Oncol. Res. 27, 1609775 (2021).

38. Chen, Y. et al. KDM4A promotes malignant progression of breast cancer by down-regulating BMP9 inducing consequent enhancement of glutamine metabolism. Cancer Cell International 24, 322 (2024).

39. Xian, F., Zhao, C., Huang, C., Bie, J. & Xu, G. The potential role of CDC20 in tumorigenesis, cancer progression and therapy: A narrative review. Medicine 102, e35038 (2023).

40. Gao, Y., Qiao, X., Liu, Z. & Zhang, W. The role of E2F2 in cancer progression and its value as a therapeutic target. Front Immunol 15, 1397303 (2024).

41. Hu, H. et al. CDCA8, a mitosis-related gene, as a prospective pan-cancer biomarker: implications for survival prognosis and oncogenic immunology. Am J Transl Res 16, 432–445 (2024).

42. Zhang, J. Z., Behrooz, A. & Ismail-Beigi, F. Regulation of glucose transport by hypoxia. Am J Kidney Dis 34, 189–202 (1999).

43. Koh, Y. W., Lee, S. J. & Park, S. Y. Differential expression and prognostic significance of GLUT1 according to histologic type of non-small-cell lung cancer and its association with volume-dependent parameters. Lung Cancer 104, 31–37 (2017).

44. Valla, M., Klæstad, E., Ytterhus, B. & Bofin, A. M. CCND1 Amplification in Breast Cancer - associations With Proliferation, Histopathological Grade, Molecular Subtype and Prognosis. J Mammary Gland Biol Neoplasia 27, 67–77 (2022).

45. Bray, F., et al. Global cancer statistics 2022: GLOBOCAN estimates of incidence and mortality worldwide for 36 cancers in 185 countries. CA: A Cancer Journal for Clinicians 74, 229–263 (2024).

46. Matsubara, D. et al. Genetic and phenotypic determinants of morphologies in 3D cultures and xenografts of lung tumor cell lines. Cancer Sci 114, 1757–1770 (2023).

47. Kinker, G. S. et al. Pan-cancer single-cell RNA-seq identifies recurring programs of cellular heterogeneity. Nature Genetics 10.1038/s41588-020-00726-6 (2020) doi:10.1038/s41588-020-00726-6.

48. Zhang, J. et al. Tahoe-100M: A Giga-Scale Single-Cell Perturbation Atlas for Context-Dependent Gene Function and Cellular Modeling. 2025.02.20.639398 Preprint at 10.1101/2025.02.20.639398 (2025).

49. DepMap, Broad. Current DepMap Release data, including CRISPR Screens, PRISM Drug Screens, Copy Number, Mutation, Expression, and Fusions. DepMap 24Q4 Public 10.25452/figshare.plus.27993248.v1 (2024).

50. Davies, A. M., Lara, P. N., Mack, P. C. & Gandara, D. R. Incorporating bortezomib into the treatment of lung cancer. Clin Cancer Res 13, s4647–4651 (2007).

51. Chua, A. D. W., Thaarun, T., Yang, H. & Lee, A. R. Y. B. Proteasome inhibitors in the treatment of nonsmall cell lung cancer: A systematic review of clinical evidence. Health Science Reports 6, e1443 (2023).

52. Knox, C. et al. DrugBank 6.0: the DrugBank Knowledgebase for 2024. Nucleic Acids Research 52, D1265–D1275 (2024).

53. Peng, B., Li, G. & Guo, Y. Prognostic significance of micropapillary and solid patterns in stage IA lung adenocarcinoma. Am J Transl Res 13, 10562–10569 (2021).

54. Yaldız, D. et al. Papillary predominant histological subtype predicts poor survival in lung adenocarcinoma. Turk Gogus Kalp Damar Cerrahisi Derg 27, 360–366 (2019).

55. Nassar, H. Carcinomas with Micropapillary Morphology: Clinical Significance and Current Concepts. Advances in Anatomic Pathology 11, 297 (2004).

56. Lee, G. et al. Clinical Impact of Minimal Micropapillary Pattern in… : The American Journal of Surgical Pathology. https://journals.lww.com/ajsp/abstract/2015/05000/clinical_impact_of_minimal_micropapillary_pattern.10.aspx.

57. Travis, W. D. et al. International Association for the Study of Lung Cancer/American Thoracic Society/European Respiratory Society International Multidisciplinary Classification of Lung Adenocarcinoma. Journal of Thoracic Oncology 6, 244–285 (2011).

58. Germain, P.-L., Lun, A., Garcia Meixide, C., Macnair, W. & Robinson, M. D. Doublet identification in single-cell sequencing data using scDblFinder. F1000Res 10, 979 (2021).

59. Parks, B. & Abdi, I. BPCells: Single Cell Counts Matrices to PCA. (2025).

60. Hao, Y. et al. Dictionary learning for integrative, multimodal and scalable single-cell analysis. Nat Biotechnol 42, 293–304 (2024).

61. Wolf, F. A., Angerer, P. & Theis, F. J. SCANPY: large-scale single-cell gene expression data analysis. Genome Biology 19, 15 (2018).

62. Leary, J. R. et al. Sub-Cluster Identification through Semi-Supervised Optimization of Rare-Cell Silhouettes (SCISSORS) in single-cell RNA-sequencing. Bioinformatics 39, btad449 (2023).

63. Chen, Y., Chen, L., Lun, A. T. L., Baldoni, P. L. & Smyth, G. K. edgeR v4: powerful differential analysis of sequencing data with expanded functionality and improved support for small counts and larger datasets. Nucleic Acids Research 53, gkaf018 (2025).

64. Korotkevich, G. et al. Fast gene set enrichment analysis. 060012 Preprint at 10.1101/060012 (2021).

65. Liberzon, A. et al. The Molecular Signatures Database (MSigDB) hallmark gene set collection. Cell Syst 1, 417–425 (2015).

66. Csardi, G. & Nepusz, T. The igraph software package for complex network research. InterJournal Complex Systems, 1695 (2006).

67. Wu, Y. et al. Neutrophil profiling illuminates anti-tumor antigen-presenting potency. Cell 187, 1422–1439.e24 (2024).

68. Kassambara, A., Kosinski, M., Biecek, P. & Fabian, S. survminer: Drawing Survival Curves using ‘ggplot2’. (2024).

69. Therneau, T. M. A Package for Survival Analysis in R. (2024).

70. Weigert, M. & Schmidt, U. Nuclei Instance Segmentation and Classification in Histopathology Images with Stardist. in 2022 IEEE International Symposium on Biomedical Imaging Challenges (ISBIC) 1–4 (2022). doi:10.1109/ISBIC56247.2022.9854534.

71. Bernstein, M. N. et al. Monkeybread: A Python toolkit for the analysis of cellular niches in single-cell resolution spatial transcriptomics data. 2023.09.14.557736 Preprint at 10.1101/2023.09.14.557736 (2023).

72. Vázquez-García, I. et al. Ovarian cancer mutational processes drive site-specific immune evasion. Nature 2022 612, 1–9 (2022).

