## Supplementary Information for "A Pan-Cancer Single-Cell Compendium of Intratumoural Heterogeneity"

**Supplementary Figures**

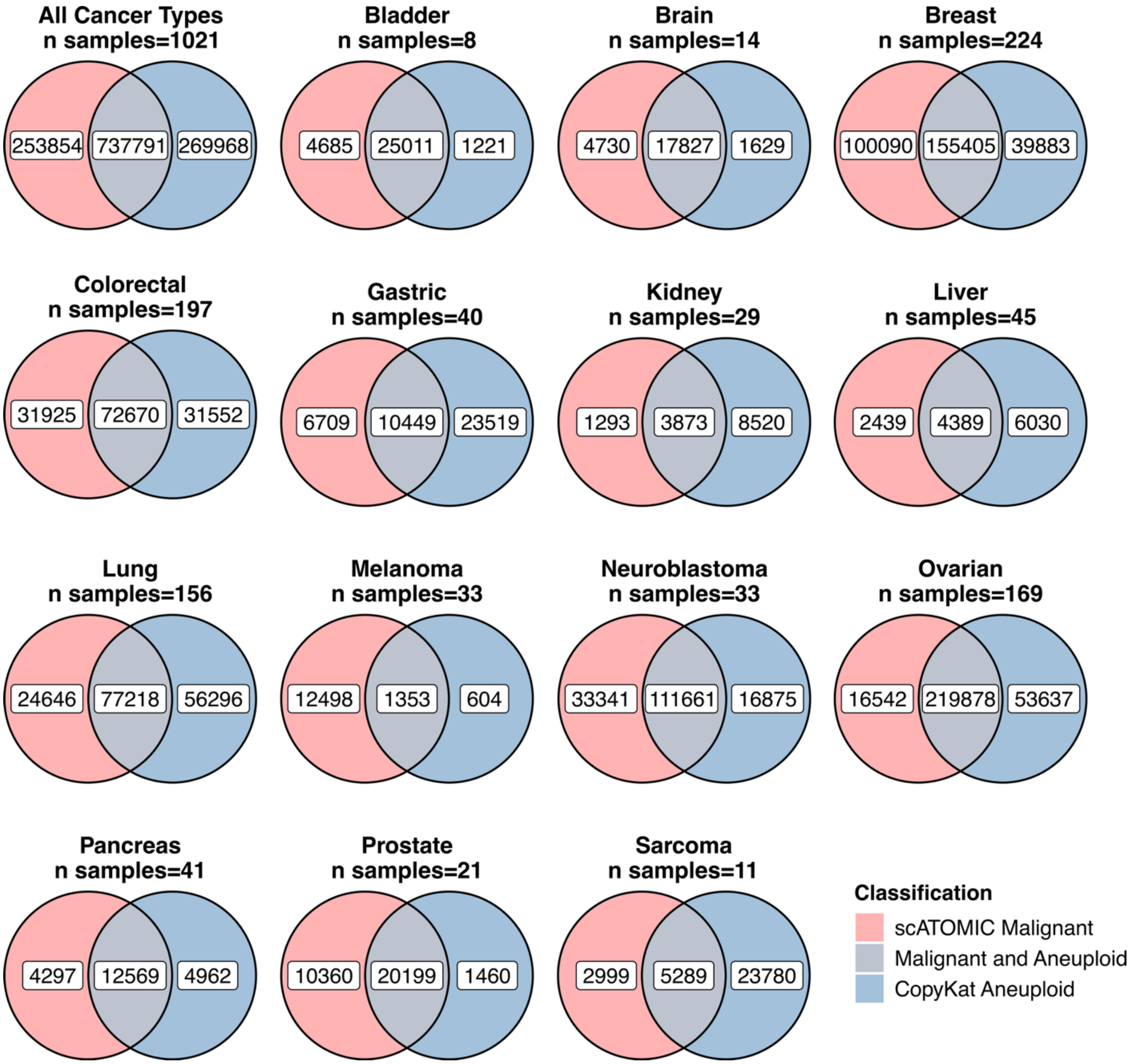


**Supplementary Figure 1: Malignant cell predictions.** Venn diagrams illustrating the overlap and discrepancies between cells classified as malignant by scATOMIC and as aneuploid by CopyKAT, shown across all samples (top left) and within individual cancer types (all other panels).

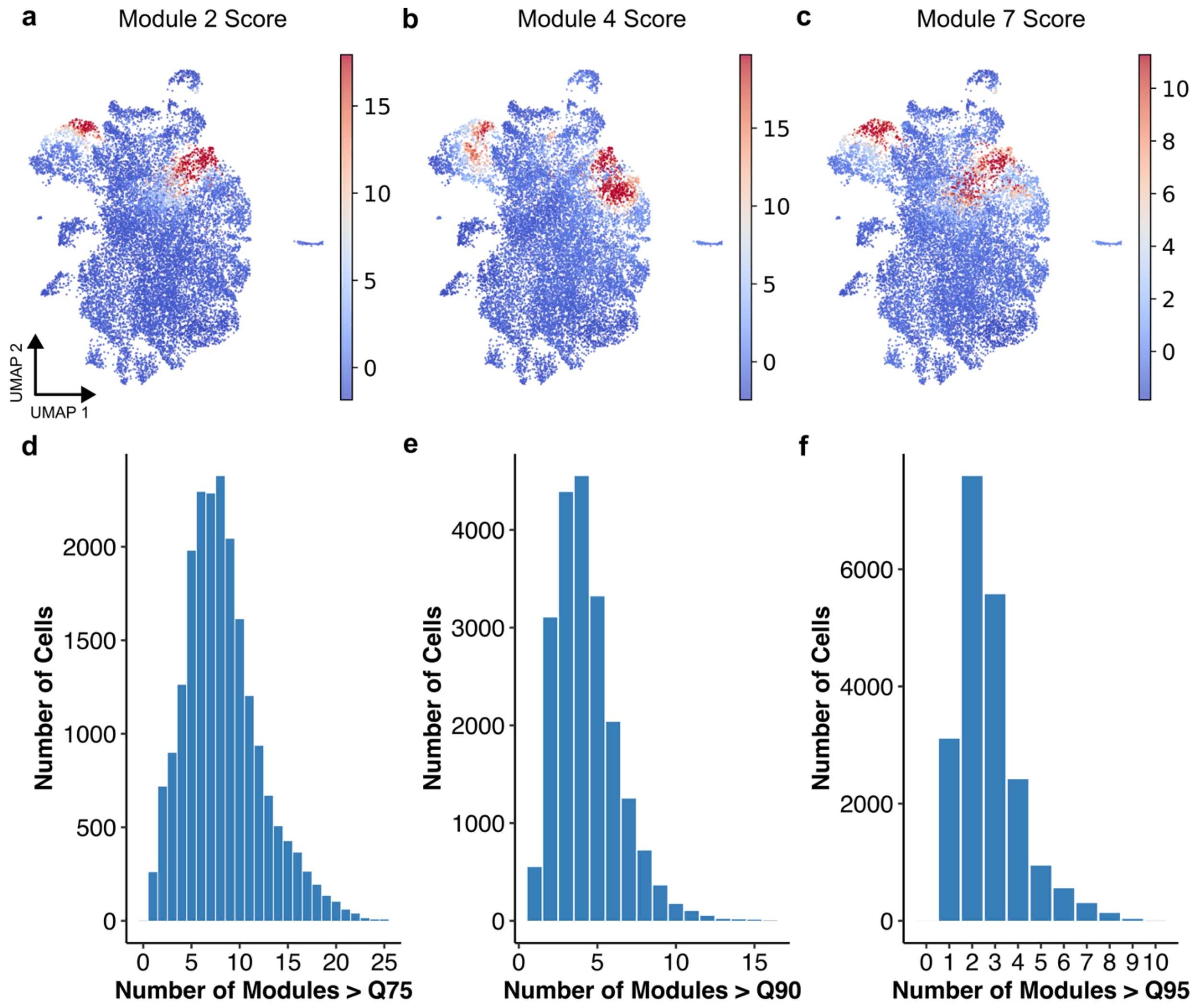


**Supplementary Figure 2: High scoring Hotspot modules across malignant cells.** Hotspot module Z-scores from Hotspot analysis in lung adenocarcinoma are shown. **a-c,** UMAP visualization of three highly correlated modules coloured by their Hotspot module Z-scores, which represent the relative expression activity of the module in each cell. **d-f,** Histograms showing the number of cells with module Z-scores above the $75^{th}$ percentile (**d**), $90^{th}$ percentile (**e**), and $95^{th}$ percentile (**f**), representing cells with high activity across multiple modules.

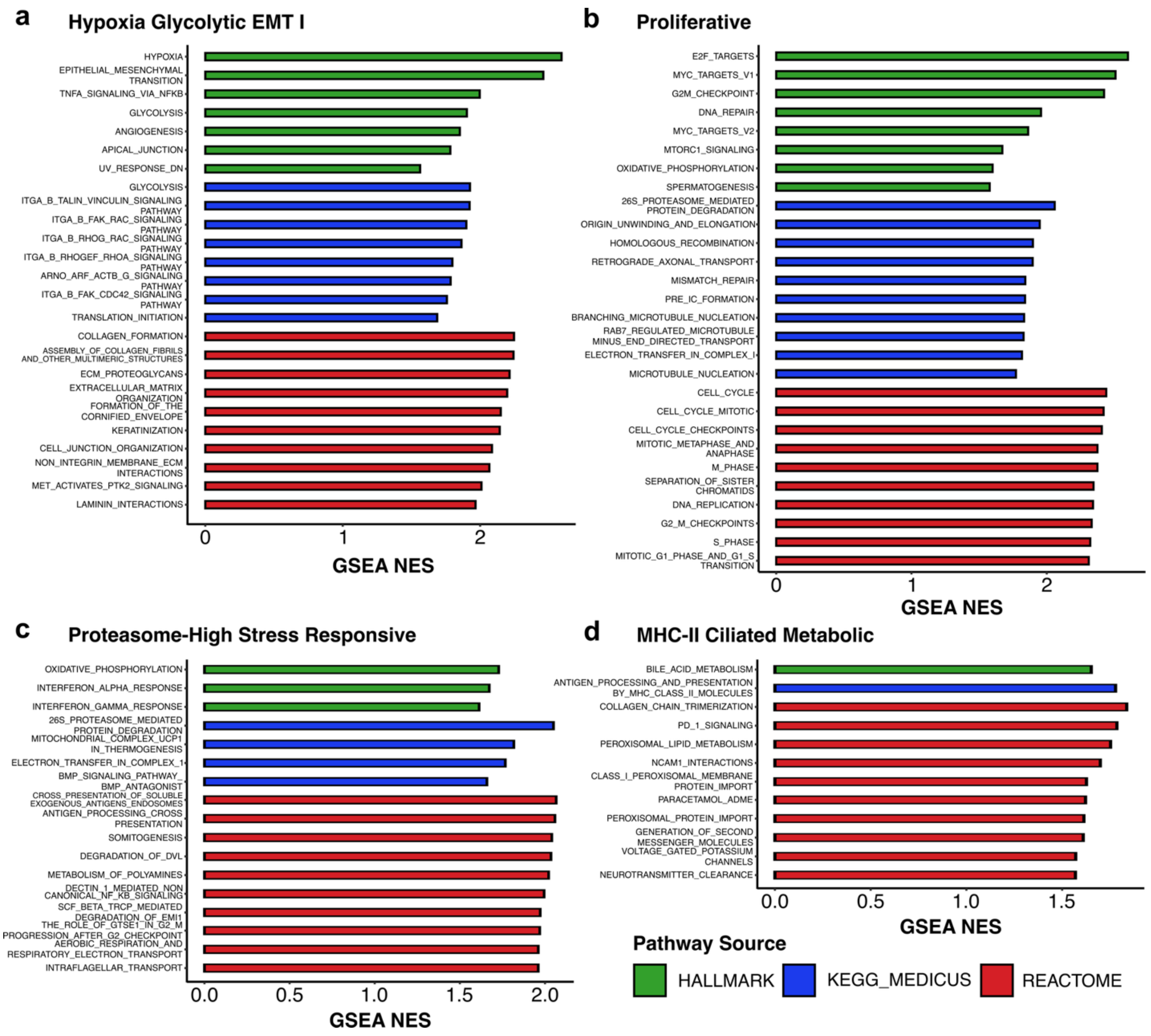


**Supplementary Figure 3: Gene set enrichment analysis in LUAD cell states. a-d**, Normalized gene set enrichment scores (GSEA NES) of top enriched pathways (P adj. < 0.05) for *Hypoxia Glycolytic EMT I* (**a**), *Proliferative* (**b**), *Proteasome-High Stress Responsive* (**c**), and *MHC-II Ciliated Metabolic* (**d**) cancer cell states. Bars are coloured by the pathway source (Hallmark gene set – green, KEGG Medicus – blue, and REACTOME – red). For each gene set database, the top enriched significant pathways are plotted, with a maximum of 10 per database.

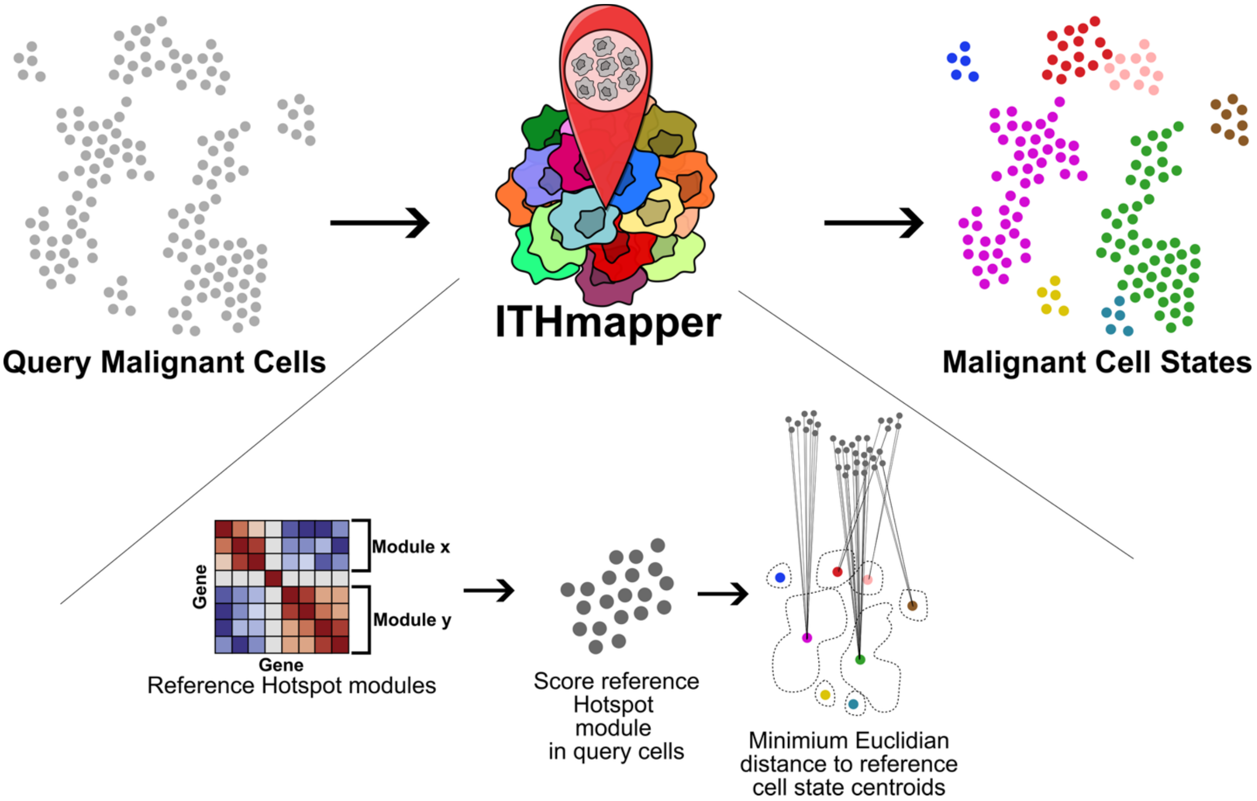


**Supplementary Figure 4: Schematic of ITHmapper.** Query malignant cells are annotated by computing Hotspot module Z-scores and projecting them into the reference module space defined by the pan-cancer atlas constructed in this study. Cell states are then assigned according to the minimum Euclidean distance to the centroid of each reference state, enabling cross-dataset mapping.

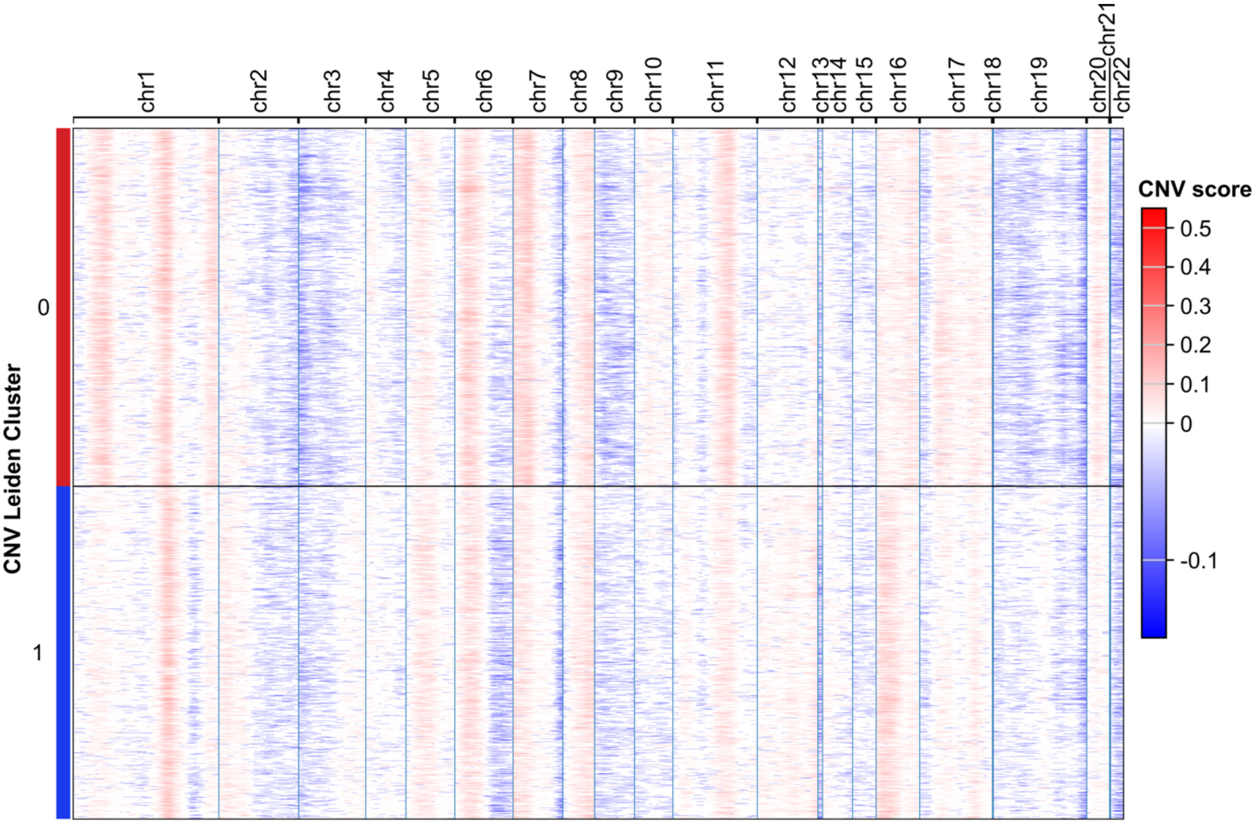


**Supplementary Figure 5: Inferred copy number variation in lung adenocarcinoma Visium HD sample of Patient 1.** Heatmap of CNV scores in malignant LUAD cells is plotted. Red and blue indicate inferred gains and losses respectively. Two Leiden clusters of CNV scores are shown on the y axis.

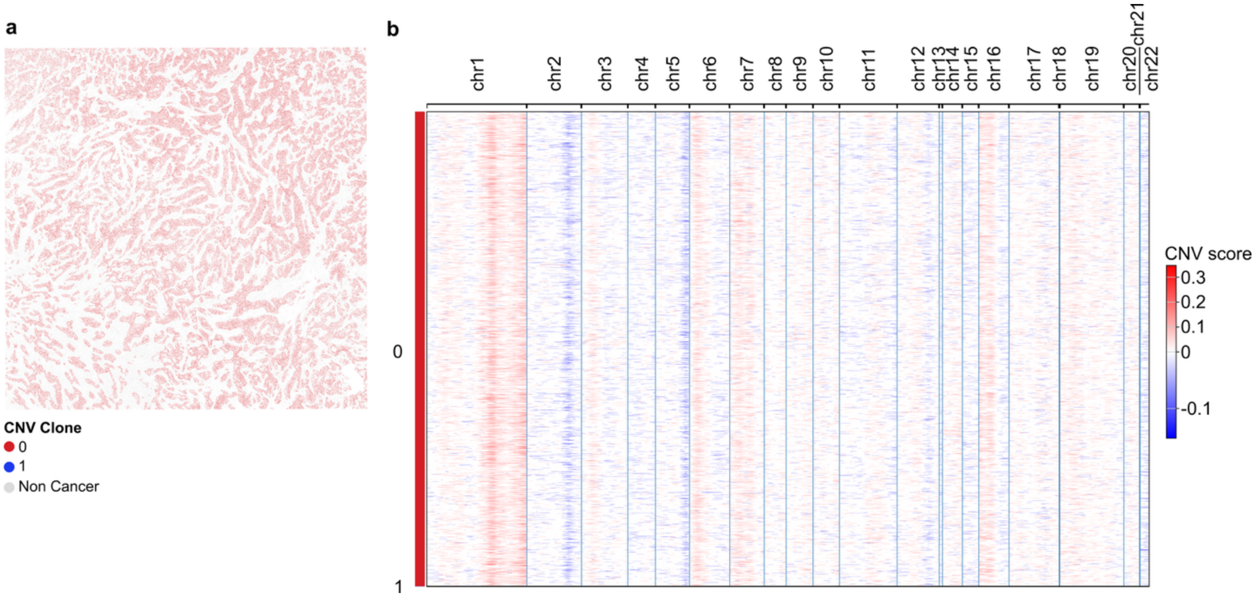


**Supplementary Figure 6: Lack of evidence for copy number variation in Visium HD breast cancer Patient 3. a**, Spatially mapped cancer cells coloured by CNV cluster. **b**, Heatmap of CNV scores in malignant BRCA with red and blue indicating inferred gains and losses respectively. Leiden clusters of CNV profiles are shown on the y-axis.
